# Testing the state-dependent model of subsecond time perception against experimental evidence

**DOI:** 10.1101/2021.12.31.474629

**Authors:** Pirathitha Ravichandran-Schmidt, Joachim Hass

## Abstract

Coordinated movements, speech, and other actions are impossible without precise timing. Computational models of interval timing are expected to provide key insights into the underlying mechanisms of timing, which are currently largely unknown. So far, existing models have only been partially replicating key experimental observations, such as the linear psychophysical law, the linear increase of the standard deviation (the scalar property or Weber’s law), and the modulation of subjective duration via dopamine. Here, we incorporate the state-dependent model for subsecond timing as proposed by Buonomano (2000) into a strongly data-driven computational network model of PFC We show that this model variant, the state-dependent PFC model, successfully encodes time up to 750 milliseconds and reproduces all key experimental observations mentioned above, including many of its details. Investigating the underlying mechanisms, we find that the representations of different intervals are based on the natural heterogeneity in the parameters of the network, leading to stereotypic responses of subsets of neurons. Furthermore, we propose a theory for the mechanism underlying subsecond timing in this model based on correlation and ablation analyses as well as mathematical analyses explaining the emergence of the scalar property and Vierordt law. The state-dependent PFC model proposed here constitutes the first data-driven model of subsecond timing in the range of hundreds of milliseconds that has been thoroughly tested against a variety of experimental data, providing an ideal starting point for further investigations of subsecond timing.

**Highlights:**

- The state-dependent model of time perception also encodes durations in the subsecond range when embedded into a data-driven network model of the prefrontal cortex.
- The model reproduces three key experimental findings of time perception: the linear psychophysical law, the scalar property (Weber’s law) and dopaminergic modulation of subjective durations.
- Many details of experimental observations can be reproduced and explained by the model without specific parameter tuning.
- A new theory for the emergence of Weber’s law and Vierordt’s law (overestimation of short durations and underestimation of long durations) is provided.
- The current variant of the state-dependent model is the first model of time perception to be thoroughly tested against a wide range of experimental evidence and can now be considered to be experimentally validated.

## 1. Introduction

The ability to estimate the passage of time is of fundamental importance for perceptual and cognitive processes [1], such as speech, music, and decision-making [2, 3]. Impairments in time perception have been reported for various psychiatric disorders, such as schizophrenia, Parkinson’s disease, Huntington’s disease, and major depression [2, 4, 5, 6, 7]. In mammalian neural circuits, time perception spans ten orders of magnitudes including millisecond timing, interval timing and circadian rhythms comprising an entire day [2, 8]. Of these ranges, specifically, the neural basis and mechanisms of estimating interval durations between hundreds of milliseconds to minutes range, remain unclear [9]. This ability, called interval timing, is involved in various processes such as foraging [10], decision-making [3], sequential motor performance [11], and associative conditioning [12, 8, 2]. Here, we focus on the subsecond range of interval timing, which is referred to as subsecond timing.

Despite the importance of telling time, the exact neural mechanisms of time perception and, in particular, for interval and subsecond timing, are not fully understood yet. Proposing and testing biologically plausible computational models is a promising way to gain an understanding of these mechanisms. A wide variety of interval timing models are currently discussed [13, 14], including the state-dependent network model [15], models of ramping activity [16], synfire chains [17], and the striatal beat model [18, 19]. Despite this wealth of modeling approaches, none of them is universally accepted as a valid theory of time perception, partly due to a lack of thorough experimental verification. Furthermore, most of the existing models deliberately abstract from many biological details, which helps to emphasize the underlying mechanism but also calls their biological relevance into question.

Here, we investigate the state-dependent network model [15], in which the timing is not encoded by a dedicated tuning mechanism but by the intrinsic neuronal and synaptic properties [20]. Specifically, Buonomano claims that the representation of time up to 400 ms is generated by time-dependent mechanisms within the network, such as short term plasticity (STP) and slow inhibitory postsynaptic potentials (IPSPs) caused by metabotropic GABA receptor (GABA_B_) synapses [15]. More recent theoretical studies suggest that the hidden states of the network, induced, for example, by STP, allow encoding any stimulus together with its temporal context and time itself in neuronal activity [20, 21, 22]. The model mimics the process of time interval estimation, where two stimuli are perceived that are set apart by a given interval and the task is to estimate the duration of that interval. The precision of such an estimate can be assessed using various measurement procedures, such as verbal estimation, reproduction, or discrimination (that is, comparison to a previously presented interval) [23, 24, 25]. While specific predictions of the state-dependent network model have been tested before [26], a thorough examination using a range of established experimental results is still missing.

In this work, we test whether the state-dependent network model constitutes a plausible neural mechanism of subsecond timing in the brain. To this end, we first implemented the model into a data-driven spiking network model of the prefrontal cortex (PFC) [27] including substantial heterogeneities in the neuron and synapse parameters. Several studies provide evidence for the central role of the prefrontal cortex in time perception in rats [28, 29, 30], monkeys [31], and humans [32, 33]. Biological constraints, and heterogeneity in particular, have previously been shown to pose a challenge to simple computational models, e.g. working memory [34]. Second, we tested whether the model could reproduce a range of established experimental findings of interval timing and can thus be validated by empirical data. Namely, we require the model to reproduce the linear psychophysical law, the scalar property, and the modulation of subjective duration by dopamine (dopamine (DA)).

The linear psychophysical law for the average duration estimates [35, 36, 23], and the scalar property for timing errors, also referred to as Weber’s law [37, 38, 39, 40, 2] have been established as fundamental properties of time perception by psychophysical experiments. The linear psychophysical law states that real-time is proportional to the averaged representation of time [35]. The linear psychophysical law is widely accepted for time perception [36, 23]. However, the slope of the linear relation is often reported to be less than one, resulting in an overestimation of short intervals and an underestimation of long intervals, a phenomenon known as Vierordt’s law[41, 42]. The slope *k* and the indifference point indifference point (IP), i.e. the duration that is essentially represented, depend on the intervals considered [43]. Studies employing intervals within the range of 200 ms-1000 ms show the IP to be between 600 ms-800 ms with the slope of *k* ≈ 0.5 [44], whereas testing intervals within the range of 400 ms-2000 ms, shifts the IP to higher values IP≈ 1400 ms with a timing slope of *k* ≈ 0.9 [45]. Another study from [46] found a slope of *k*≈ 0.7 for visual stimuli and *k* ≈ 1.1 for auditory stimuli for short intervals (400 ms-600 ms) and a slope of around *k* ≈ 0.6 for longer intervals (2000 ms-3000 ms). Although some studies suggest that IP can be calculated by the arithmetic mean of the applied intervals [45, 43, 47], other studies do not support this hypothesis [48, 49, 43].

The scalar property or Weber law refers to the observation that the timing error, i.e., the standard deviation of the duration estimates *σ*, increases linearly with the duration of the interval. To quantify this relation, the Weber fraction can be used, dividing the standard deviation of the duration estimate by its mean. A constant Weber fraction over a range of durations is an indicator of Weber’s law. Although observed in numerous studies, the scalar property is not universal but seems to hold for intervals of around 200 ms [50, 51, 52, 53] up to 2000 ms [50, 54] in humans, with typical Weber fractions of around 0.05 - 0.13 [55, 50]. For shorter intervals, the Weber fraction increases, indicating a sublinear (e.g., square root) scaling of timing errors, while for longer intervals, a superlinear increase in timing errors and an increase in Weber fraction is reported in different species (pigeons [53], rats [56] and humans [50, 54]). Using information-theoretical methods, [57] suggest that these different scaling regimes of timing errors may indicate different mechanisms underlying time perception. In particular, the scalar property is explained by so-called variance-based timing in this framework, i.e., using the temporal information contained in the timing errors itself, which increase over time, and thus can be used to decode time by means of a pure diffusion process.

While the linear psychophysical law and the scalar property constitute static properties of time perception, several factors such as emotions and mood [39], attention [58] and stimulus properties [59, 60, 61] have been described to modulate time perception. A prominent pharmacological modulator of time perception is dopamine (DA) [2, 62]. Specifically, studies suggest that of the two main types of DA receptors, D1 and D2, the latter has a high potential to modify time perception [63, 64]. Numerous studies report that decreasing dopaminergic activity, e.g., by dopaminergic antagonists, can slow down the internal clock leading to an underestimation of timing intervals, while increasing levels of DA, e.g., by dopaminergic agonists, speeds up the internal clock [2, 62, 64]. Although most of these studies have been conducted for intervals in the seconds to minutes range [2, 65], similar effects have also been found for shorter intervals, investigating, for example, dopaminergic drugs in healthy humans [66, 67, 68, 69] and mice [64] and dopamine-related conditions such as ADHD [70, 71, 72].

We find that the state-dependent network model reproduces all three experimental findings outlined above within a limited range of interval durations when implemented within the PFC network model [27]. Analyzing the underlying mechanisms, we further found that the network can be dissected into neuron pools, each of which represents a specific interval. In particular, a subset of neurons within a pool respond maximally to a specific stimulus representative of a time interval. As confirmed by ablation of the model components, this duration-specific response is mainly driven by STP and GABA_B_. Furthermore, limitations for longer interval durations and higher Poisson noise are examined to test the robustness of the model. Finally, we propose a theoretical account for the generation of subsecond timing following Weber’s law within this model that also hints at possible generalizations beyond the particular model presented here and links violations of Weber’s law with the occurrence of Vierordt’s law. This study can be seen as an experimental validation of the state-dependent network model. To the best of our knowledge, this model is the first that has been tested against such a wide range of experimental results.

## 2. Results

In order to test its timing properties in a more realistic setting, we incorporated the state-dependent model proposed by Buonomano [15] into a biologically detailed PFC model [27] (see Methods for details). Briefly, the model consists of 1000 simplified adaptive exponential (simpAdEx) integrateand-fire neurons [73], which are subdivided into superficial (L2/3) and deep (L5) cortical layers and into pyramidal neurons and various types of interneurons. Neurons are randomly connected by conductance-based double exponential functions of the types AMPA, NMDA, and GABA_A_ equipped with three types of dynamics STP, namely facilitation, depression, and a combined version of both [74, 75]. All parameters were randomly drawn from experimentally validated distributions, and synaptic connectivity was based on pairwise connection probabilities from the literature [27]. Neurons are driven by a constant background current (*I*_back_) that differs between pyramidal cells and interneurons. To implement the state-dependent model, a slow GABA_B_ [76] current was included into the PFC network model, compensated by increasing *I*_back_ by a factor of 1.4. A comparison of the spike statistics with and without GABA_B_ can be found in Suppl. Fig. A.1. Except for a decreased coefficient of variation (CV), the statistics did not change by adding GABA_B_ to the model.

Layer 2/3

All neurons were stimulated at the beginning and end of an interval with a short step current (*I*_*s*_ = 220 pA, duration 10 ms), cf. Fig. 1. The first stimulus was applied to the network at 1500 ms to ensure that the network has reached steady state. For convenience, the starting point of the first stimulus will be defined as *t*_0_ = 0 ms. After simulation, the number of spikes of each neuron within a time window (∆*t*_*w*_ = 25 ms) that ends at 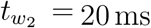 after the second stimulation (see Fig. 16A in the Methods section), were extracted. These numbers of spikes are referred to as states and are assumed to contain information about the interval duration between the two stimuli. Raster plots of all neurons for exemplary inter-stimulus intervals are shown in Suppl. Fig. A.2.

**Figure 1:**
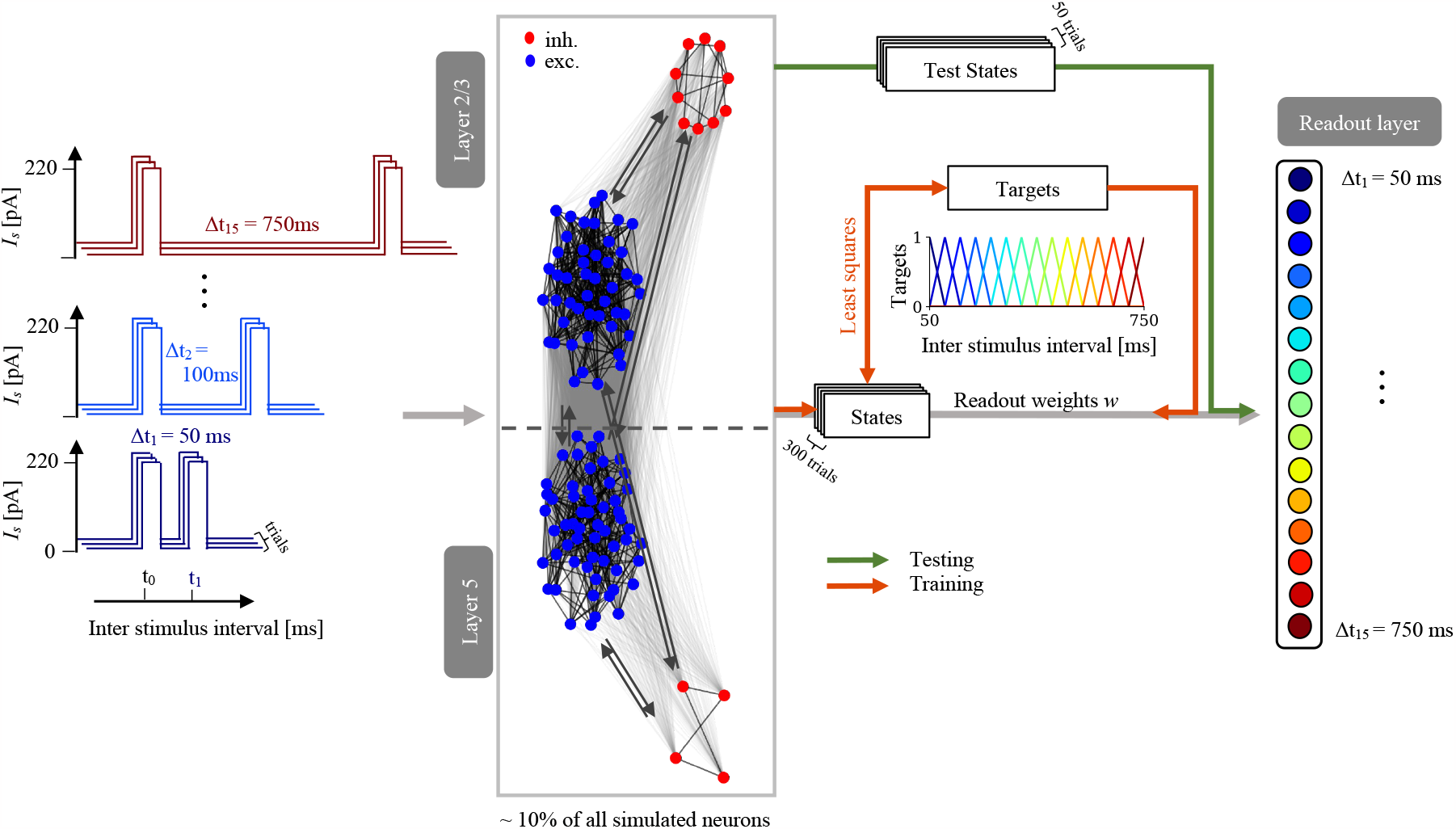
Illustration of the interval timing model. The PFC network (center gray box) with layer 2/3 and layer 5 neurons (excitatory: blue, inhibitory: red, connectivities: gray arrows) is stimulated in the beginning and the end of several inter-stimulus intervals (color-coded from 50 ms to 750 ms). Spike trains are extracted within a window around the second stimulus to compute network states and to train weights of the readout layer via least squares to predict the estimated inter-stimulus intervals ∆*t*_test_.

To predict interval durations ∆*t*_test_, a readout layer was added to the model with the number of output units *N*_*r*_ matching the number of interval durations presented during training, cf. color-coded readout units in Fig. 1. Each output unit is then trained with the least squares method [77, 78] (see Methods for details) to be active for the respective time interval and to be inactive for all other time intervals. We emphasize that this training is not intended to mimic learning processes in the brain. Rather, the readout layer is used as an ideal observer that extracts as much temporal information out of the state-dependent network as possible, as in the original publication [15]. We trained the weights of the readout units with various inter-stimulus intervals (∆*t*_train_ = 50 - 750 ms) in 50 ms steps and tested in 25 ms steps. Normalized outputs of trained readout units for different test intervals ∆*t*_test_ are shown in Fig. 2A. Each readout unit has a bell-shaped tuning curve peaking at its respective interval and being nonzero at neighboring time points, allowing the model to generalize to intervals it has not seen during training. For intervals longer than 300 ms, the tuning curves become broader.

**Figure 2:**
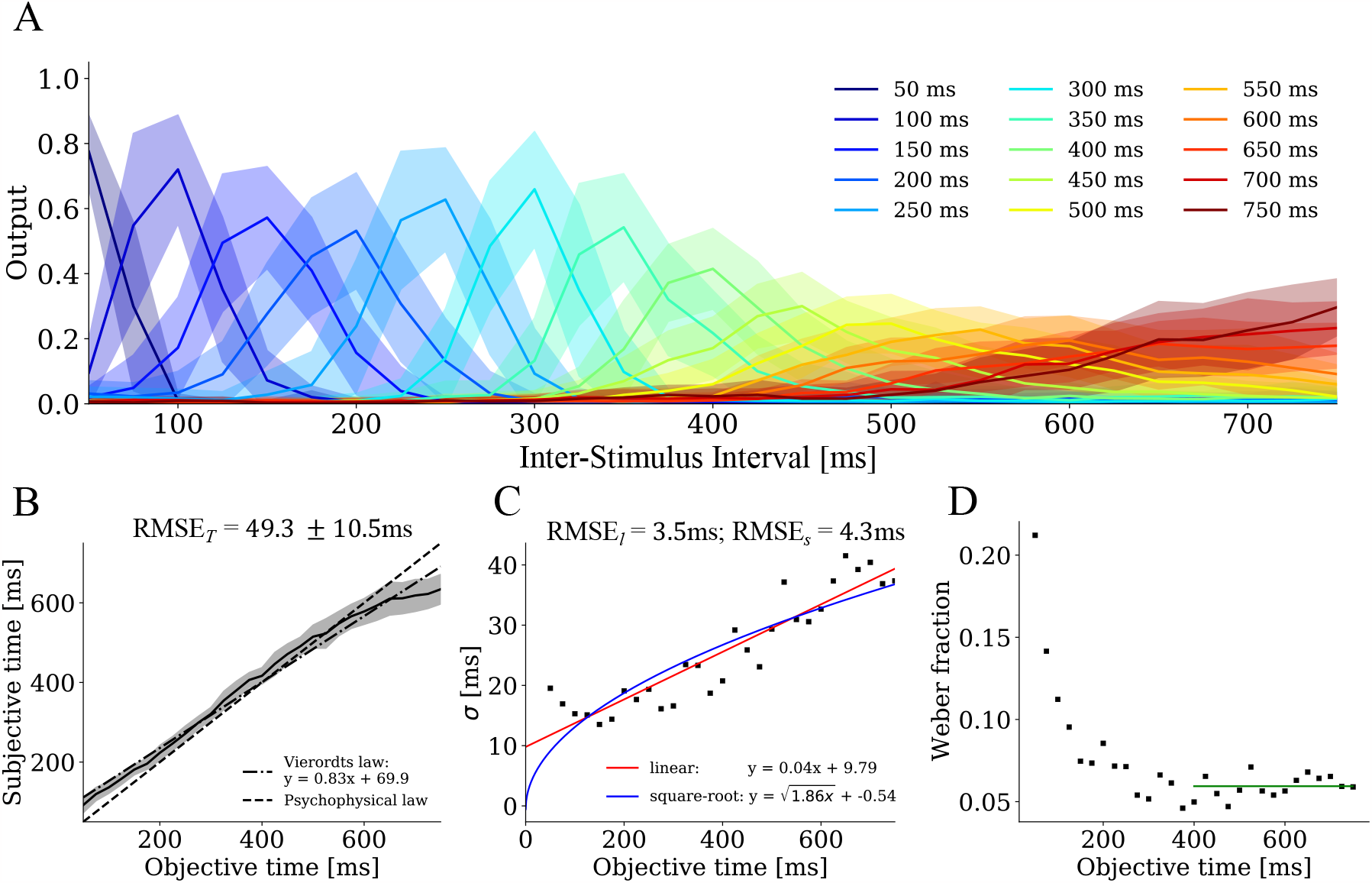
Psychophysical law and scalar property. **A** Tuning curves of the readout units (color-coded) with standard deviation for different test intervals (x-axis). **B** Averaged subjective time (over 50 trials) as a function of test interval durations together with the standard deviations (shaded region), the linear fit (dashed-dotted-line) and the objective time (dashed black line). **C** Standard deviations of the estimated time as a function of test interval durations, fitted to a linear (red curve) and to a square root function (blue curve). **D** The trial-averaged Weber fraction as a function of test interval durations. The horizontal line (in green) depicts the average value between (400 - 750 ms).

### Testing the linear psychophysical law and the scalar property

To test the linearity of the psychophysical law, we calculated the average subjective time by multiplying the output values of the readout units by their corresponding ∆*t*_test_ and adding all readout units. The estimated time is well described by a straight line (Fig 2A), with an overestimation of shorter intervals and an underestimation of longer intervals, in agreement with Vierordt’s law. A linear fit yields slopes *a* = 0.8 ±0.1 and offsets *b* = 69.9 ±16.0. The slopes *a* are within the range of experimental observations [46, 44] and IP of the averaged estimated times can be found at IP = 530 ms, also in line with experimental studies [49].

To test the scalar property or Weber’s law, we computed the standard deviations of the estimated time over 50 trials, as shown in Fig. 2C, which is fitted with a linear function (in red) and for comparison with a square root function (in blue). We also computed the trial-averaged Weber fraction, restricted to the range of 400 - 750 ms (green line in Fig. 2D) to control for the fact that constant errors (the offset of the line) dominate at shorter intervals, causing the Weber fraction to decrease (generalized Weber’s law [53]). The Weber fraction of 0.059 matches the experimental observations [50, 79]. Computing the readout weights with a ridge regression instead of linear least squares yields almost identical results, cf. Suppl. Fig. A.3.

Both properties, the linear psychophysical law and the scalar property, were further tested by averaging over several sets of neuronal and synaptic parameters, each of which is randomly drawn from experimentally validated distributions [27]. For ten parameter sets, the psychophysical law is well-fitted by a linear function and a slope of 0.85 ±0.05, which is significantly smaller than 1.0 (one sample t-test: *t* (499) = − 74.8, *p <* 0.001). The IP of the averaged estimated time is 545.5 ±23.3 ms.

Regarding the scalar property, Fig. 2C suggests that it is not trivial to determine whether the standard deviation follows a linear or a square root function, i.e., whether the scalar property holds true. To answer this question, we averaged the standard deviations over simulations for the same ten parameter sets, but with a larger number of ∆*t*_test_ (10 instead of 25 ms steps, Fig. 3). Up to interval durations of 500 ms, the data are well approximated by a linear function. However, at longer ∆*t*_test_, the standard deviations saturate to a constant value. Overall, the data can be approximated by a piecewise linear function of the form

**Figure 3:**
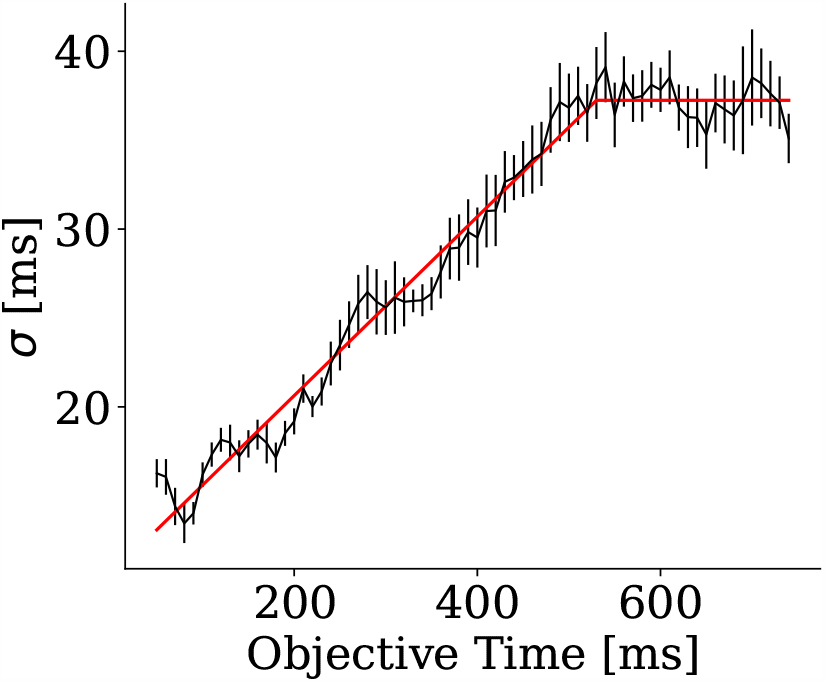
Assessing the scalar property. Standard deviations of the estimated time as a function of test interval durations averaged across ten different parameter sets. The red line shows the best fit to a piecewise linear function, saturating for ∆*t*_test_ above a threshold (see the text for details). The ten different parameter sets were only used for this figure.

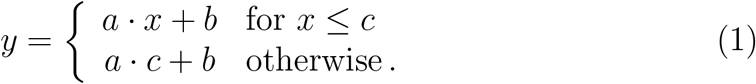

The parameters of the piecewise linear fit *a, b*, and *c* are shown in Table 1.

**Table 1:** Fit parameters for the scalar property over ten parameter sets with ∆*t*_test_ = 10 ms.

| Fit type | fit parameters |  |  |
| --- | --- | --- | --- |
| | $a$ | $b$ | $c$ |
| piecewise linear | $0.06 \pm 0.02$ | $10.2 \pm 2.5$ | $530.0 \pm 89.5$ |
| linear | $0.04 \pm 0.01$ | $13.9 \pm 1.6$ | - |
| square root | $1.9 \pm 0.7$ | $2.6 \pm 2.9$ | - |

### Dopaminergic modulation

To examine the third result, the modulation of subjective duration by the D2 dopamine receptor, we simulated a transient change in the activation of the D2 receptor by changing the neuronal and synaptic parameters by the values summarized in Table 5, which reflect the experimentally observed effects of D2 *in vitro* (see Methods for details). These changes were only applied for the test phase, while for training, the same parameters were used as before. The changes in Table 5 were considered to be activation of 100 % D2 (D2 agonist), while the original parameters reflected activation of 0 %. Inverting the sign of all changes in Table 5 simulated − 100 % deactivation of D2 (D2 antagonist), so 0 % represented a baseline activation level that can be increased or decreased.

The full range of altered time estimates for DA modulation from −100 % to 100 % and for ∆*t*_test_ of 50 - 750 ms are presented in Suppl. Fig. A.4. As both agonistic and antagonistic modulations show boundary effects for DA modulations *>* ±50 % and for the longer and shorter intervals (Suppl. Fig. A.4), we only considered intervals within a range of 200 ms - 600 ms and DA modulations up to± 50 %, see Fig. 4A. The shift of the activation of the output neurons for increasing dopaminergic modulation is shown in the Suppl. Fig. A.5 for the 400 ms-encoding output neuron.

**Figure 4:**
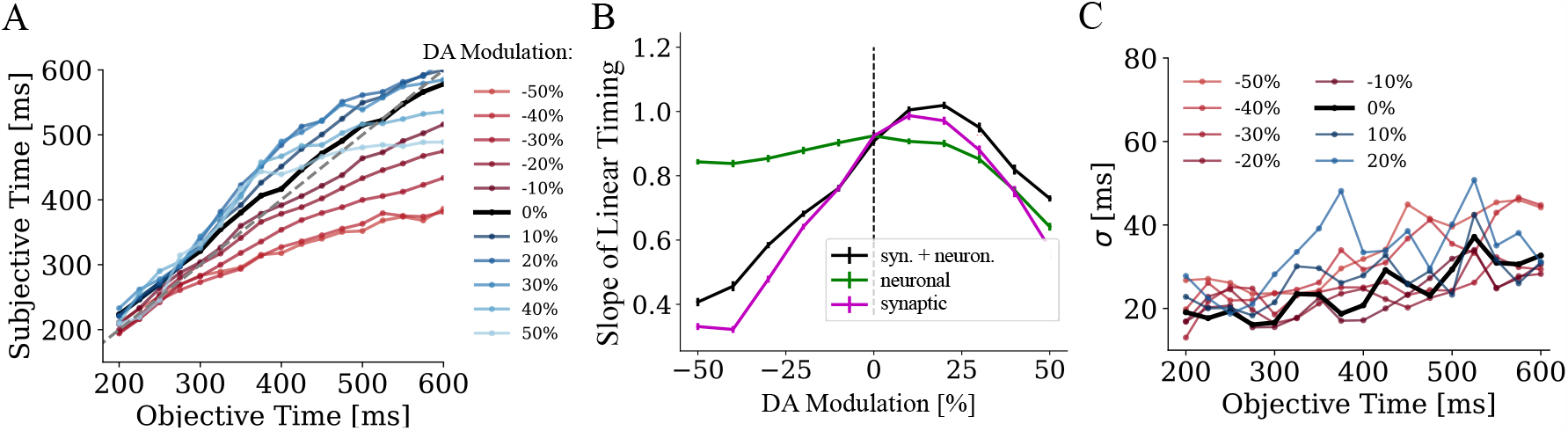
Effects of dopaminergic modulation. **A** Subjective duration as a function of test interval durations between 200 ms - 600 ms without modulation (black curve), for agonistic (blue curves) and antagonistic (red curves) D2 modulation. **B** The slope of the linear psychophysical law for each modulation of synaptic and neuronal parameters together (black curve), for synaptic modulation alone (magenta curve) and for neuronal modulation alone (green curve). The dashed vertical line indicates the point without modulation. **C** Standard deviations of the estimated time as a function of test interval durations for D2 modulation between −50 % and 20 %.

Within this limited range of durations and modulation levels, subjective time shortens for higher antagonistic modulations (Fig. 4A, from darker to brighter red colors) and lengthens for agonistic modulation (Fig. 4A, from darker to brighter blue colors), consistent with experimental results [2, 62]. In Fig. 4B (black curve), we show the slope of a linear fit to the psychophysical law for each level of D2 modulation. The slopes reach a maximum around 20 % (one-way ANOVA over 50 trials, *F*(10, 539) = 299.3, *p <* 0.001) and drop again for higher values. To further assess the mechanism of subjective time modulation of D2, we restricted modulation to synaptic parameters (Fig. 4B, magenta curve) and neuronal parameters (Fig. 4B, green curve). For synaptic modulation, the slopes basically follow the same curve as with both modulations together, whereas the neuronal modulation alone shows only an effect at strong agonistic modulation, where it further decreases the slope.

To assess a potential dopaminergic modulation of timing errors (Fig. 4C), we also fitted a linear function to the standard deviations within the range of modulations where subjective duration linearly increases (− 50 % to 20 %), but observed systematic changes neither in the slope (linear regression: slope = −0.03, *p* = 0.20, *R*^2^ = 0.26) nor in the intercept (linear regression: slope = 9.62, *p* = 0.16, *R*^2^ = 0.30) of Weber’s law. The standard deviations and the slopes over the full range of durations and modulations are shown in Suppl. Fig. A.6.

So far, we have simulated acute D2 modulation in the test trials, while leaving the training trials unaffected by dopamine. If we retrain the readout weights under the influence of D2 modulation, we can counteract the changing speed of the internal clock, as we tested on − 30 % agonistic and antagonistic modulation. Fig. 5 shows the psychophysical law and Weber’s law for agonistic dopaminergic modulation of 30 % (red) for the antagonistic dopaminergic modulation of −30 % (blue), both of which are almost identical to the unmodulated simulation from Fig. 2B (black).

**Figure 5:**
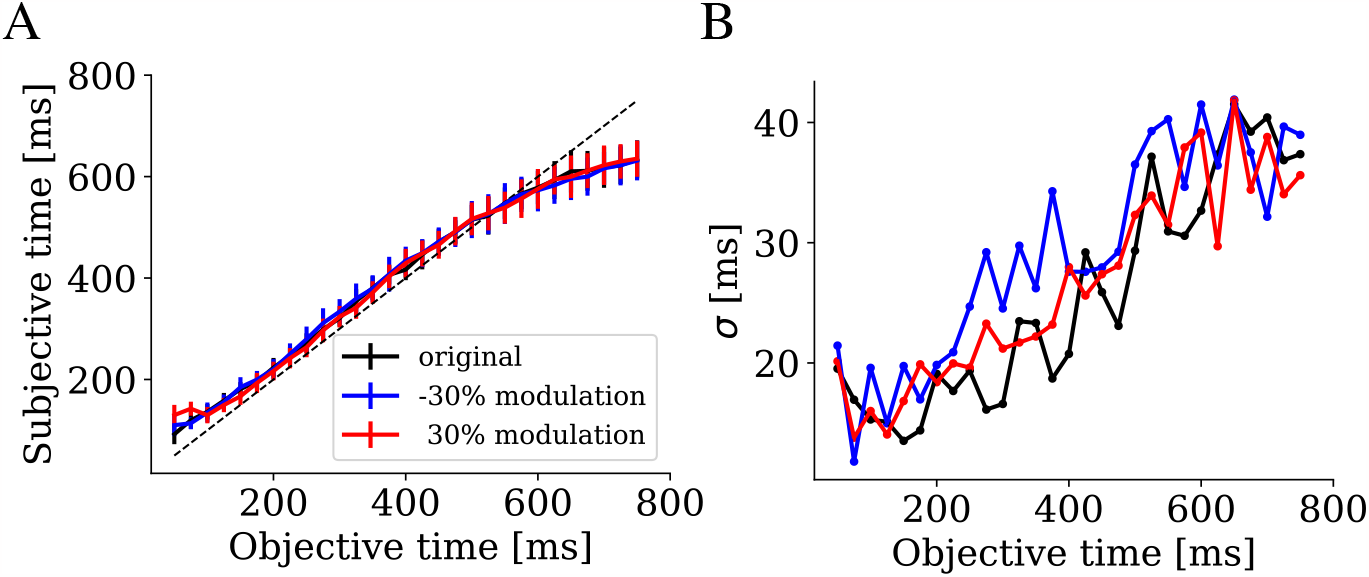
Retraining after mild dopaminergic modulation. Estimated times (panel **A**) and the corresponding standard deviations (panel **B**) as a function of test interval durations for agonistic (30 %, red) and antagonistic modulation (− 30 %, blue) after retraining. The estimated times and the standard deviations without modulation were re-plotted from Fig. 2 in black for comparison.

### Limitations of the model

To identify potential limitations of the model, we considered longer intervals and higher noise levels. When we trained the readout network for longer interval durations up to 2000 ms and tested within the same range, we found that the time perception was less accurate. In particular, the slope of the psychophysical law is much smaller compared to the shorter range, implying a more pronounced Vierordt’s law (slope = 0.7 ±0.1) with the IP shifted towards longer intervals (IP = 852.9 ±186.3 ms), cf. Fig. 6A. While the slope still matches experimentally observed values 0.5 − 1.1 [44, 45, 46], the IP for a similar range 400 ms-2000 ms is higher at around ≈ 1400 ms in experiments [45]. Fitting parameters for the standard deviations are *a* = 0.3, *b* = −3.9 and *c* = 615.0, cf. Fig. 6B.

**Figure 6:**
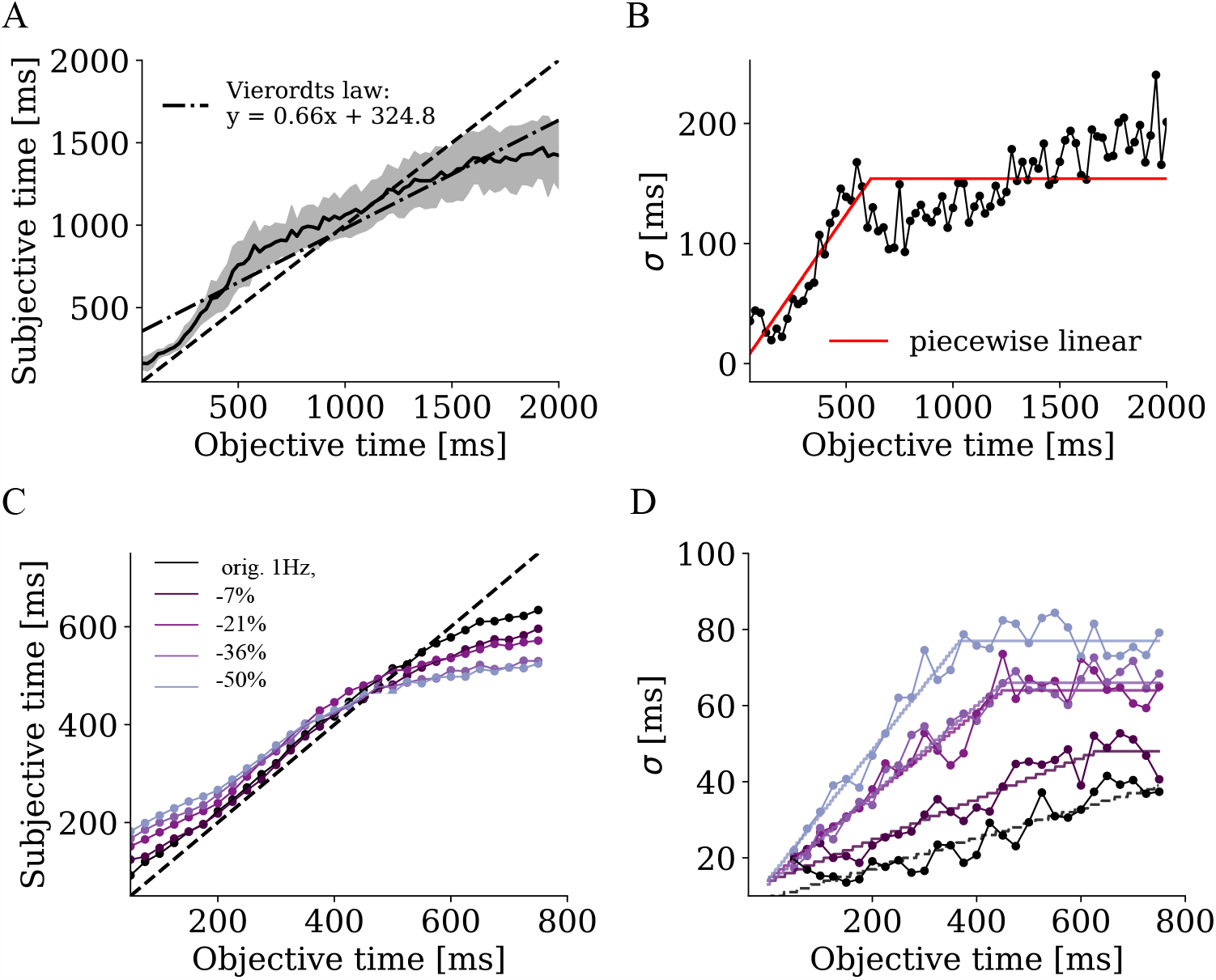
Testing for limitations of timing in the model. Estimated time (panels **A, C**) and standard deviations (panels **B, D**) as a function of test interval durations for longer durations up to 2000 ms (panels **A, B**) and for higher noise levels (panels **C, D**). Lighter colors in panels **C** and **D** represent higher noise levels (see legend in **C**). The corresponding Poisson weights and the fit parameters are shown in Table 2.

Regarding the effect of noise levels, we originally connected *N*_Poisson_ = 10 Poisson neurons with a firing rate of 1 Hz to the network. However, a typical cortical neuron has approximately 1000-10 000 connections with other neurons [80, 81] and can thus be subject to much higher noise levels. To investigate whether subsecond timing still works in this case, we increased the firing rates of the 10 Poisson neurons to *f* =1000 Hz and compared interval timing at different noise levels by increasing the Poisson neuron weights *w*_Poisson_ and simultaneously decreasing the background current to avoid overexcitation and keep the standard deviation of the subthreshold membrane potential (see Suppl. Fig. A.7) constant, cf. Table 2 and Fig. 6C and D.

**Table 2:** Parameters for different noise levels with 1000 Hz Poisson neurons.

| $I_{\text{back red.}}$<br>[%] | $w_{\text{Poisson}}$ | Psychophysical law | | Scalar property | | |
| --- | --- | --- | --- | --- | --- | --- |
|  |  | slopes | IP<br>[ms] | piecewise linear fit |  |  |
|  |  |  |  | a | b | c |
| 0 (1 Hz) | 0.5 | $0.8 \pm 0.1$ | 530.0 | 0.04 | 9.8 | 910.4 |
| -7 | 0.003 | $0.7 \pm 0.1$ | 472.0 | 0.05 | 14.2 | 625.0 |
| -21 | 0.005 | $0.7 \pm 0.1$ | 512.5 | 0.1 | 14.7 | 454.2 |
| -36 | 0.007 | $0.6 \pm 0.1$ | 477.7 | 0.12 | 13.5 | 452.5 |
| -50 | 0.010 | $0.5 \pm 0.1$ | 458.7 | 0.17 | 14.7 | 370.6 |

We find a less accurate timing for increasing noise levels: Timing errors increase more quickly over time and Vierordt’s law becomes more pronounced as noise levels increase; see Table 2. The piecewise linear function provides a good fit for the standard deviations (Table 2) for all noise levels, but the cut-off value *c* is shifted towards shorter intervals for higher noise levels (see Fig. 6D).

### Mechanisms of subsecond timing

Given that interval timing, especially Weber’s law, works within the statedependent PFC model at least for limited noise levels and intervals up to 750 ms, we next aimed to understand the underlying mechanisms of timing within this model. First, we performed ablation analyses, i.e., systematically disabled specific components of the network to determine which of them are critical for which aspect of timing in the model. Second, we performed correlation studies to determine whether there are pools of neurons that encode specific intervals and, if so, how they differ from neurons that are predictive of other intervals. Finally, we studied a simplified model to explain the origin of the scalar property and Vierordt’s law.

### Ablation analyses

In this section, we systematically remove some of the components of the model to analyze which are critical for our results. We focus on three main ingredients: synaptic processes with long time constants, heterogeneity of neuronal and synaptic parameters, and irregular background activity (induced by constant background currents).

Regarding synaptic dynamics, there are three elements with long time constants that might influence the estimation of time: NMDA currents, GABA_B_ currents, and STP of each of the synaptic currents. To test their specific influence on time estimation, we conducted ablation analyses by removing each of these synaptic processes, adjusting the background current and synaptic weights to compensate for missing input, and retraining the readout layer. The exact adjustments can be found in Suppl. Table A.1. The parameters of Vierordt’s law (slope and IP) are shown in Suppl. Table A.3, while the fit parameters for the piecewise linear fit for Weber’s law are presented in Suppl. Table A.4.

Removing NMDA synapses had no effect on estimated times, see Fig. 7A and D in magenta, Suppl. Table A.3 and Suppl. Table A.4. In contrast, when removing STP, time estimation gets worse reflected in a much more pronounced Vierordt’s law and increased standard deviation compared to the original state-dependent PFC model, see Suppl. Table A.3, Suppl. Table A.4 and Fig. 7A,D in red. This effect is even more pronounced when GABA_B_ is removed, such that inhibition is limited to GABA_A_ with much faster time constants, see Fig. 7A,D in blue. To test the combined effect of GABA_B_, STP and NMDA, we removed all three of them in a further simulation, which yielded a slope of 0.35±0.04 for time estimation and an even higher standard deviation and decreased slope *a* compared to ablating GABA_B_ alone, see Fig. 7A,D in cyan.

**Figure 7:**
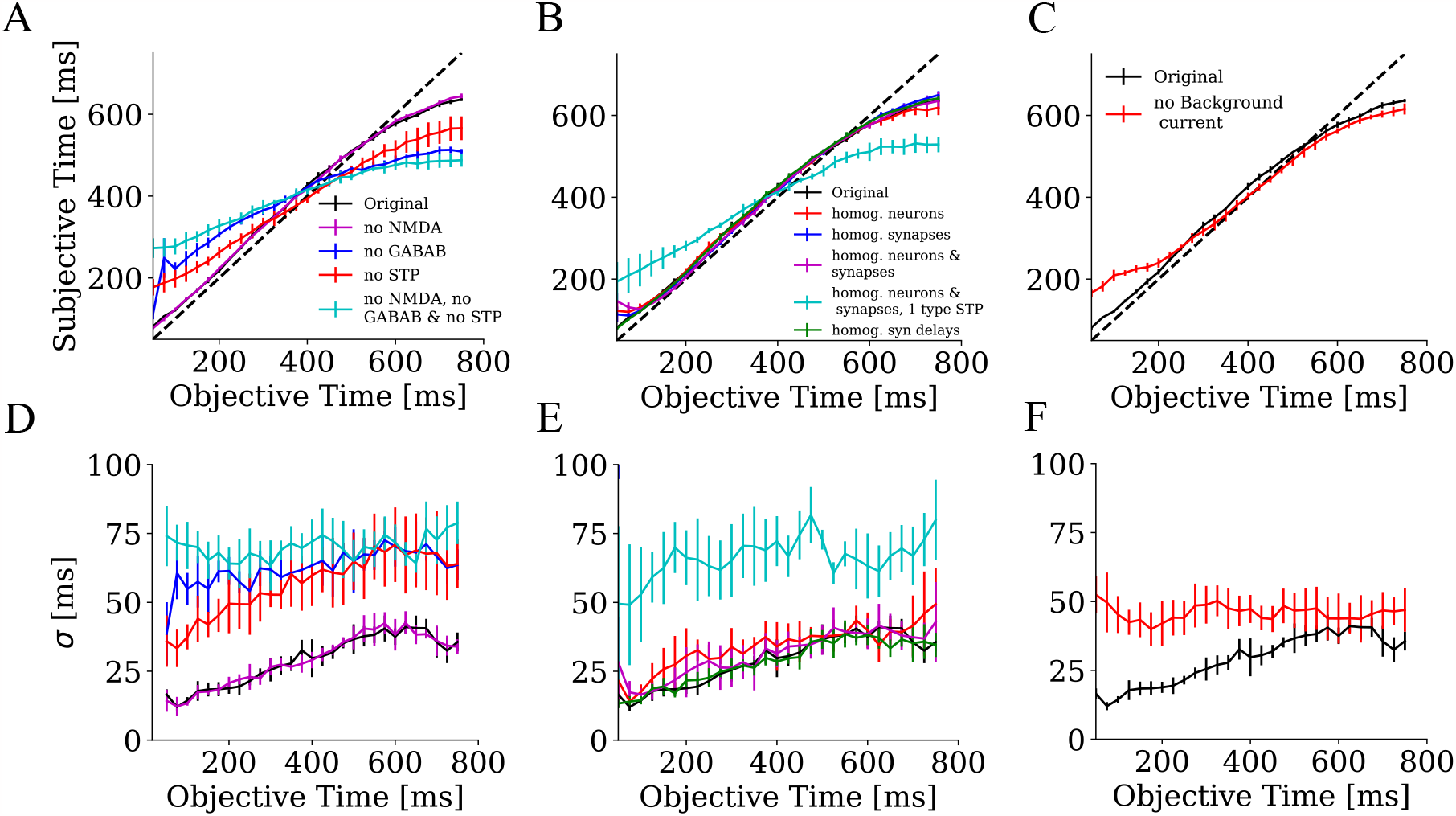
Ablation studies. Estimated durations (panels **A-C**) and their standard deviations *σ* (panels **D-F**) averaged over five parameter sets as a function of test interval durations for different ablations. **A, D** Removing synaptic processes, namely NMDA (magenta curves) and GABA_B_ (blue curves) currents, short-term plasticity (STP, red curves) and the combination of all three (cyan curves). **B, E** Removing heterogeneity of the neuronal parameters (red curves), of the synaptic parameters (blue curves), and of both (magenta curves), additional reduction to one type of STP within each pair of neuron types (light blue curves) and removing the heterogeneity of synaptic delays only (green curves). **C, F** Removing the background current within the PFC model (red curves). In all panels, the original results from Fig. 2B and C are re-plotted in black. The corresponding fit parameters are shown in Suppl. Table A.3 and A.4.

In the next step, we tested the role of heterogeneity in neuron and synapse parameters. In the original simulations, these parameters are randomly drawn from distributions that are specific for each neuron and synapse type [27]. Here, we used homogeneous parameters within each neuron type. To this end, we first removed the heterogeneity within each neuron type for the neuronal parameters, second, for the synaptic parameters connecting each pair of neuron types, and finally, for both parameter sets. For all three ablations, we observed similar slopes of the linear psychophysical law and a slightly shifted IP only for homogeneous neuronal and synaptic parameters, see Suppl. Table A.3. We then further reduced heterogeneity by using only one single type of STP for each pair of neuron types (see Methods for details). This manipulation resulted in a much more pronounced Vierordt’s law with a slope of around 0.53 ±0.08 and a reduced IP as well as higher standard deviations with larger offset *b*, see the cyan line in Fig. 7B and E and Suppl. Table A.3 and Suppl. Table A.4. Finally, using homogeneous synaptic delays (which are also randomly drawn in the original simulations) for all synapses does not affect timing, see green line in Fig. 7B and E and Suppl. Table A.3 and Suppl. Table A.4.

In order to determine whether the background activation of neurons in between two subsequent stimulations might be responsible for the linear timing results, we removed the background current to all neurons, such that no pronounced activity could be seen between intervals. Removing the background current only impaired timing accuracy up to 200 ms (Suppl. Table A.3 and Fig. 7C) and disrupted Weber’s law to a negative slope near zero (Suppl. Table A.4 and Fig. 7F). Furthermore, we showed in the previous section that increasing noise levels up to 1000 Hz yielded larger timing errors and gradually shifted the cut-off value *c* of the piecewise linear fit of the scalar property towards shorter intervals (see Table 2 and Fig. 6C and D).

### Interval-encoding pools

To understand the key features of interval discrimination within training, we classified each neuron in the network as encoding a specific interval according to the readout neuron with the highest readout weights, following the procedure in [15]. All neurons encoding the same interval form a so-called interval-encoding pools (IEPs). Only neurons with positive readout weights of above 0.01 are considered in the following. In this way, we find interval-encoding pools (pool size:*μ* = 14.3, *σ* = 4.6, min = 4, max = 23), of which 571 out of 1000 neurons were not assigned to an interval-encoding pool, as the highest readout weights are too small. After assigning neurons to pools, the pools were sorted from shorter to longer intervals (see Fig. 8A). Within each interval-encoding pool, the neurons were additionally sorted from low to high weights.

**Figure 8:**
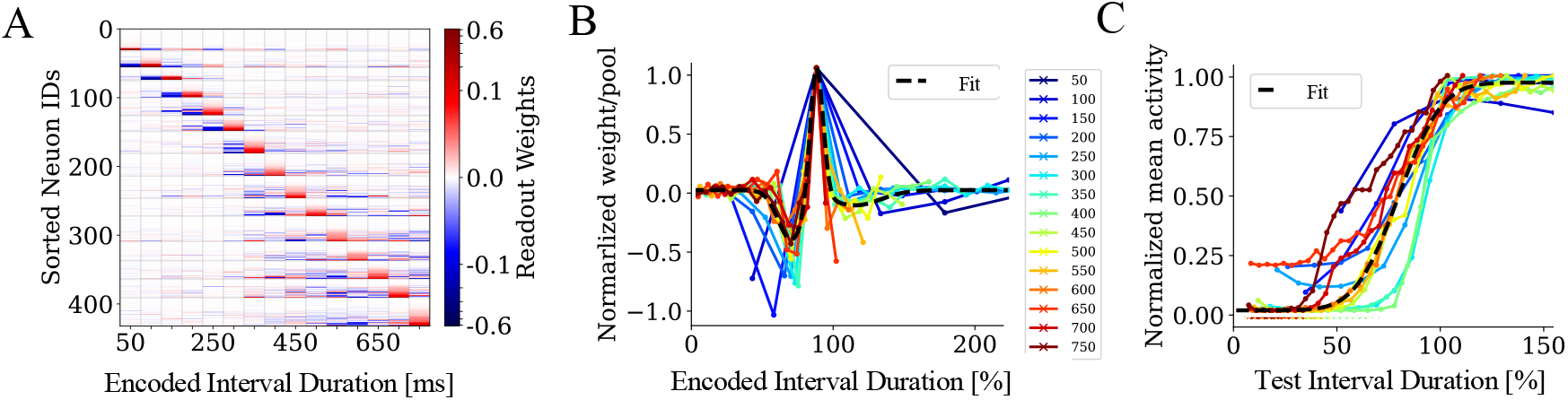
Readout weights and average firing rates within interval-encoding pools. **A**. Neurons are associated with readout units based on the largest weight *w* compared to the weights of other readout units. The weight matrix is then first sorted by the association to readout units, and within each group, sorted by their strongest weights to the respective readout unit. The horizontal lines show the borders of the pools of neurons, and the vertical lines separate the encoded intervals. The color scale follows a log_10_-scale. **B** Normalized averaged readout weights of neurons within a pool as a function of the normalized encoded interval durations of each pool (color-coded by the readout intervals). The dashed line is the best fit to a sum of two Gaussians of all curves. **C** The averaged states over 50 trials within a pool normalized for each test interval normalized by the encoded duration of each pool color-coded by the readout intervals, see legend in **B**. The dashed line represents the best fit of the stereotypic firing rate profiles to a sigmoid function (dashed line).

Calculating the normalized average weight distribution from the neurons in each group to their respective readout neuron, we see stereotypic weight profiles which we term *temporal receptive fields*: Weights peak shortly before the time that the respective pool represents and suppress contributions from both earlier and later pools asymmetric (Fig. 8B). Furthermore, we compared the states of neurons in each pool in response to the second stimulus and also found a stereotypic firing rate profile for most pools: the normalized firing rate increases until 100 % of the elapsed time for the respective pool is reached and saturates for all longer intervals, see Fig. 8C. Taken together, these results indicate that each pool translates the time elapsed relative to the interval it represents into a similar firing rate code and also transfers this code to its respective readout neuron using a stereotypic weight profile.

### What makes a pool interval specific?

Having confirmed the functional relevance of the interval-sensitive pools of neurons, this section is concerned with the mechanism that underlies this sensitivity. To this end, we compared synaptic currents at different points in time between different pools, as well as a number of neuronal and synaptic parameters.

First, we examine the sum of all synaptic currents that affect each group, separated for excitatory and inhibitory currents, as shown in Figs. 9A and B. As expected, all synaptic currents peak during the first stimulus and again at the second stimulus.

**Figure 9:**
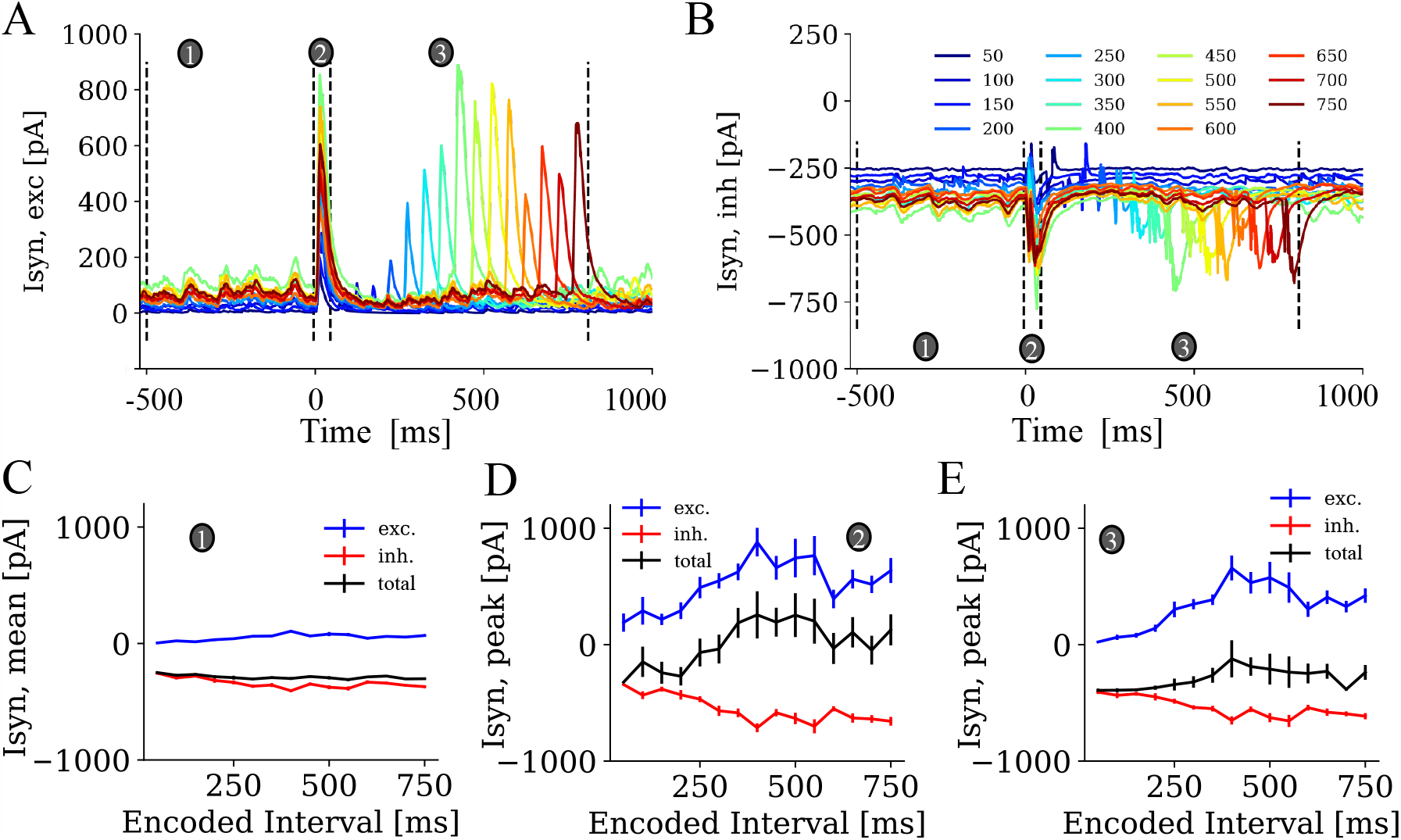
Synaptic currents within the pools. Average excitatory (panel **A**) and inhibitory (panel **B**) synaptic currents of neurons within each group as a function of simulated time. The three circles and the dashed lines mark three time ranges for which the synaptic currents are compared across pools, namely before stimulation (*t* = − 500 ms to *t* =− 5 ms, marked as 1, panel **C**), during first stimulation (*t* = − 5 ms to *t* = 45 ms, marked as 2, panel **D**), and during the second stimulation (*t* = − 5 ms + ∆*t* to *t* =− 5 ms + ∆*t* + 50 ms marked as 3, panel **E**). For each case, average excitatory (blue curves), inhibitory (red curves), and overall currents (black curves) are shown for each pool as a function of the duration encoded in each pool.

The absolute amplitude of synaptic currents was compared for the trained intervals before the first stimulus (Fig. 9C) by averaging over the range depicted with the circle (1), then the peak values were computed in response to the first stimulus (Fig. 9D, (2) and in response to the second stimulus (Fig. 9E, (3) between pools. Both excitatory and inhibitory currents increase in pools encoding longer intervals ∆*t* in all three regimes (Fig. 9C, D, and E). This linear effect is most pronounced for ∆*t* up to 400 ms, but was still statistically verified for the whole range of ∆*t*, see Table 3. Also, the total current (the sum of excitatory and inhibitory currents) results in a statistically significant increase across IEPs (see Table 3).

**Table 3:**
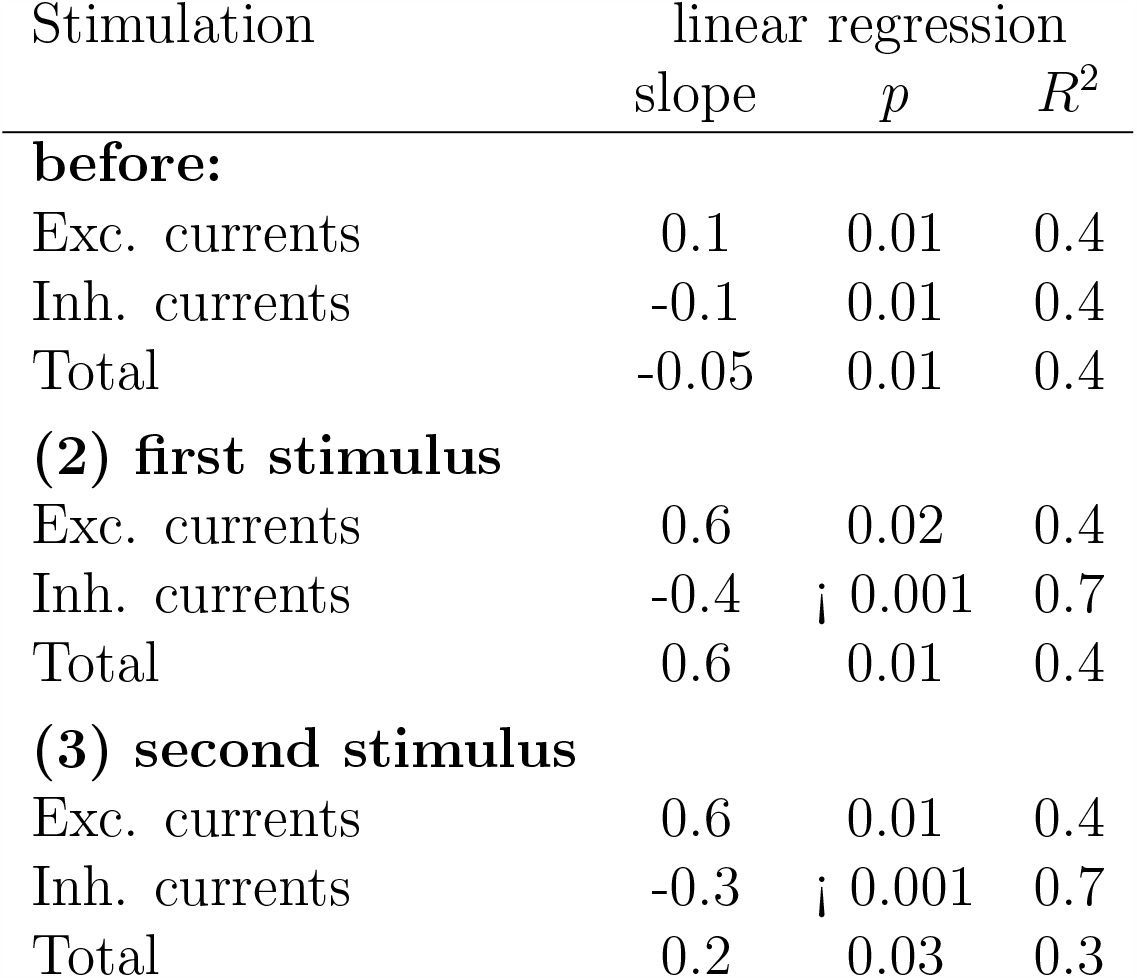
Synaptic currents within IEP averaged before stimulation (*t* = −500 ms to *t* = −5 ms), after the first stimulation (*t* = −5 ms to *t* = 45 ms) and after the second stimulation (*t* = −5 ms + ∆*t* to *t* = −5 ms + ∆*t* + 20 ms) for all encoding intervals.

To determine the role of neuronal excitability in encoding different intervals, we calculated the average rheobase for each group (minimal current to make each neuron fire, Fig. 10A, see Methods for details) and subtracted this value from the average total synaptic current within the range *t* = − 500 ms to *t* = − 5 ms for each neuron, cf. the black line in Fig. 9C. While the rheobase (slope = 0.02, *R*^2^ = 0.13, *p* = 0.2) does not differ between pools, both the total synaptic current (Table 3 before stimulation, total) and the mean (neuron-wise) difference between rheobase and total synaptic current, see Fig. see Fig. 10B significantly differ over the whole interval range between 50 and 750 ms (linear regression: slope = 0.07 and *R*^2^ = 0.8, *p <* 0.01).

**Figure 10:**
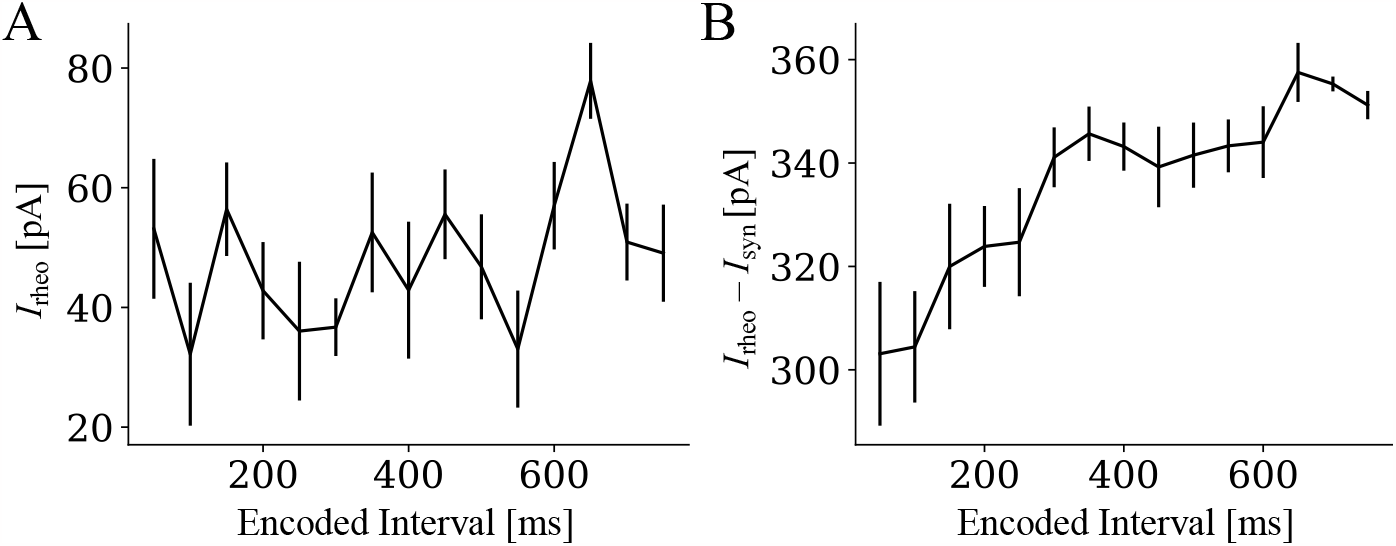
Excitability of interval-encoding pools. **A** Averaged rheobase (see Methods for details) and **B** the difference of the rheobase to the total synaptic current within each pool before stimulation as a function of the duration encoded in each pool, cf. black line in Fig. 9C.

Finally, we examine whether differences in synaptic currents between pools are determined by differences in neuronal and synaptic parameters (Fig. 11). Indeed, the summed synaptic weights onto all neurons within each IEPs (Fig. 11A) and the number of synaptic inputs onto each pool (11B) increase in pools encoding longer intervals, for both inhibitory and excitatory synaptic connections as well as the sum of the two. We also found an increase in the averaged synaptic delays for higher intervals for excitatory and both synapses (Fig. 11C) and an increase in the averaged membrane time constants *τ*_*mem*_ across pools for intervals up to 400 ms (linear regression: slope = 0.03, *p* = 0.008, *R*^2^ = 0.72)(Fig. 11F). Furthermore, the facilitating time constants within the excitatory (Fig. 11D) and inhibitory neurons (Fig. 11E) show an increase for intervals up to ∼ 400 ms. At the same time, the depression time constants *τ*_ref_ are decreased up to ∼ 400 ms for both excitatory neurons (Fig. 11D) and (Fig. 11E) for inhibitory neurons. The linear regression parameters for each of these cases are shown in Suppl. Table A.2.

**Figure 11:**
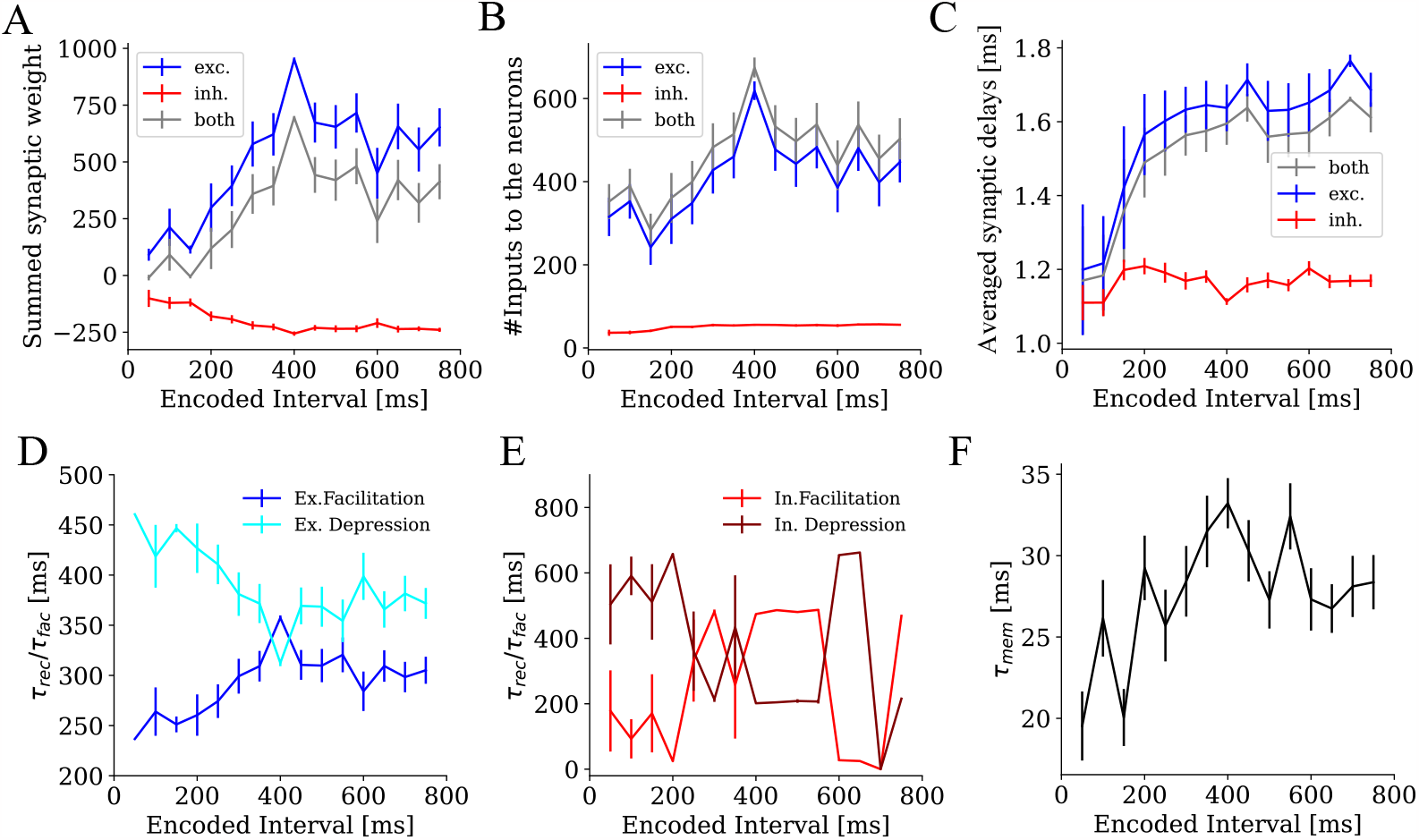
Neuronal and synaptic properties within IEPs, as a function of the duration encoded in each pool. **A** Averaged synaptic weights onto neurons within IEPs for positive weights only (blue curves), negative weights only (red curves), and all weights (grey curves). **B** Same as in **A** for the number of inputs and **C** for the average synaptic delays. **D** Averaged time constants of short-term facilitation (*τ*_*fac*_) and depression (*τ*_*rec*_) for excitatory neurons within IEPs. **E** Same as **D** for inhibitory neurons. **F** Averaged membrane time constants *τ*_mem_ for both neuron types within each pool.

### Origin of the scalar property

Here, we discuss how the scalar property of the timing errors may arise in the state-dependent model. To this end, we first recall how the duration estimate *T*_est_(*t*) is computed from the network and the output neurons in mathematical terms (Fig. 12). We then compute the mean and the standard deviation of this estimate for a special case to show that the scalar property arises from a) the scalar invariance of the firing rates of the neurons in the IEPs and b) the coupling between mean and standard deviation of the firing rates by means of the binomial distribution. In the Appendix, we illustrate how this derivation can be generalized under the assumption that the tuning curves of the output neurons follow the same shape for all intervals. Finally, we explain how the scalar invariance of the firing rates in the neuron pools of the network comes about using a minimal model that captures the main features of the full network.

**Figure 12:**
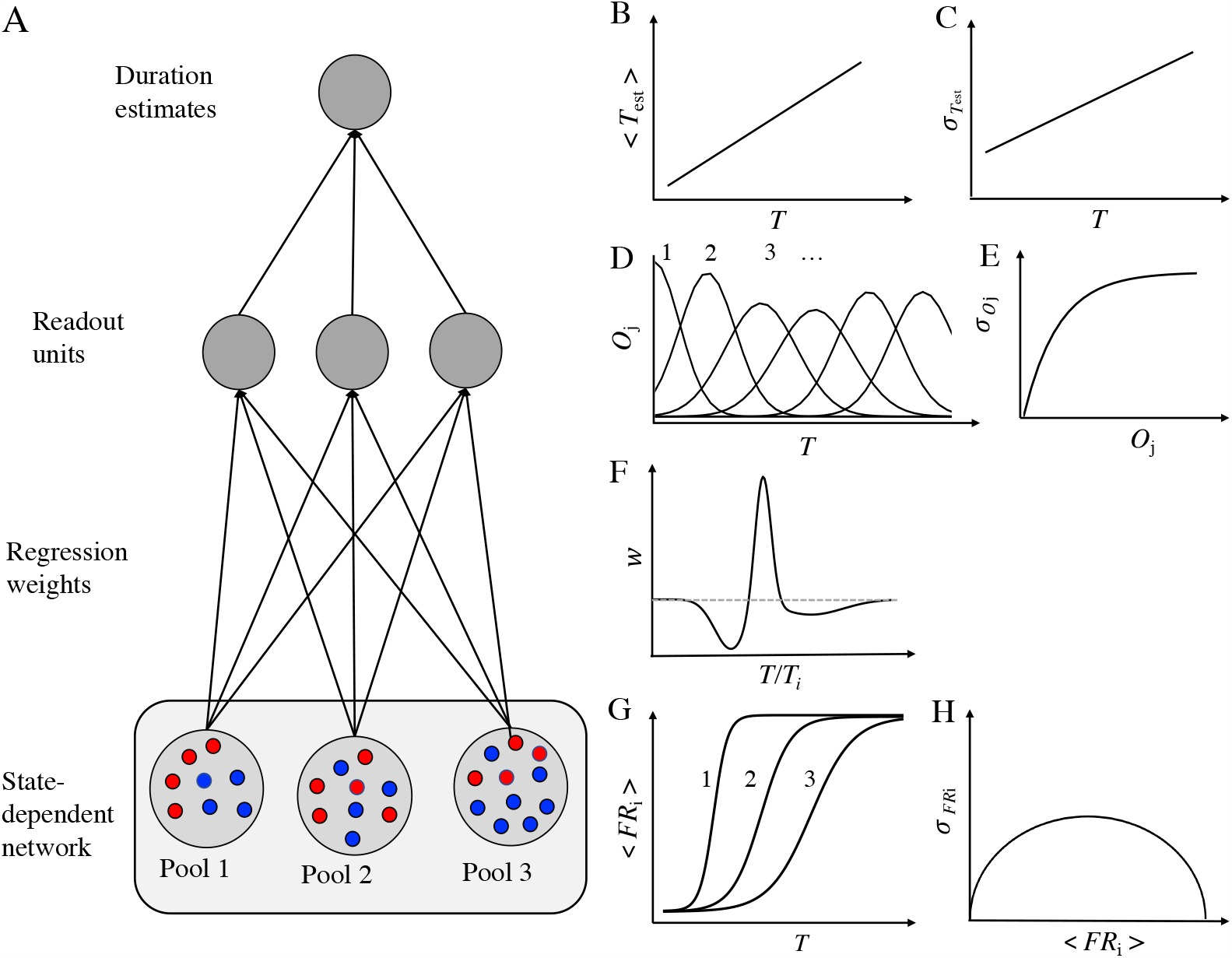
Overview on the derivation of the scalar property. **A** Translating the activity within interval-encoding pools (bottom, blue: excitatory, and red: inhibitory) into timing estimates and their standard deviations (top). **B**,**C** Multiplying the readout units with their corresponding time duration and summing up over all readout units leads to the estimated times and the scalar property. **D** Tuning curveshaped outputs are generated within the readout neurons with a defined relation between mean and standard deviation (panel **E**). **F** Synaptic weights from the pools to the output neurons, forming a stereotypic *temporal receptive field*. **G** Firing probabilities peaking at one close to the encoded time of each of the pools. **H** Relation of mean and standard deviations of the firing rates, following from the binomial distribution.

As summarized in Fig. 12A, durations are estimated in the model using the state-dependent network, divided into *N* IEPs, and the same number of output neurons, each of which encoding an interval duration *T*_*i*_. The activity levels of the output neurons *O*_*i*_(*t*) are multiplied by the respective duration *T*_*i*_ and summed to form the duration estimate *T*_est_(*t*). Furthermore, the outputs *O*_*i*_(*t*) arise from the firing rate states FR_*i*_(*t*) of each interval-encoding pool of the network, multiplied by the output weights *w*_*ij*_. Thus,

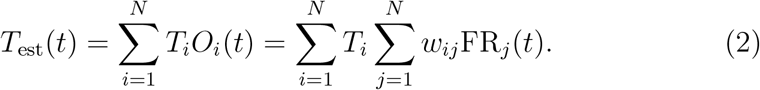

To highlight the main features of the model that give rise to the scalar property, we first restrict ourselves to estimate durations that are identical to the ones that are encoded in a single interval-encoding pool (*t* = *T*_*i*_) and reduce the model to contain only the one output neuron associated with this interval-encoding pool. Under these assumptions, Eq. 2 reduces to

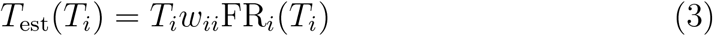

The scalar property requires that 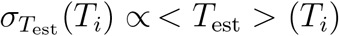, where 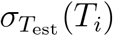 and *< T*_est_ *>* (*T*_*i*_) are the mean and standard deviation of *T*_est_ at time *T*_*i*_, respectively. As *T*_*i*_ and *w*_*ii*_ are constant, it follows 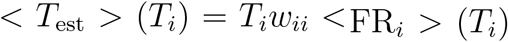 and 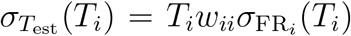. Two features of the model allow us to directly relate these two moments: First, as seen in Fig. 8C, the firing rate curves FR_*i*_(*t*) follow a stereotypic and time-invariant shape for all IEPs *i*. Thus, one can write FR_*i*_(*t*) = FR(*t/T*_*i*_). In particular, for *t* = *T*_*i*_, the firing rates are FR_*i*_(*T*_*i*_) = FR(*T*_*i*_*/T*_*i*_) = FR(1), which is constant across pools. Second, note that each pool consists of *N*_*i*_ neurons, each with firing probability *p*_*i*_(*t*) at a given time *t*. If each neuron is assumed to fire not more than one spike in response to the second stimulus (which is true for the vast majority of neurons within all IEPs), the firing rate (i.e., the number of firing neurons) follows the binomial distribution (Fig. 12H). This distribution describes the number of successes (here: spikes) in a sequence of *N* random experiments (here: neurons), where each success has the same probability *p*. The mean firing rate *< FR*_*i*_ *>*= *N*_*i*_*p*_*i*_ of the pool *i* and its standard deviation 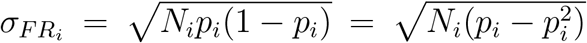 are known from this distribution. Using the first relation, the mean firing rate at time *T*_*i*_ in pool *i* can be expressed as FR_*i*_(*T*_*i*_) = FR(1) = *N*_*i*_*p*∗, where *p*∗ is the (constant) peak firing probability of all pools at the time *T*_*i*_ that pool represents.

From the above considerations, and by setting the weight *w*_*ii*_ to 1/(*N*_*i*_*p*∗), such that *< T*_est_ *>* (*T*_*i*_) = *T*_*i*_, it follows that

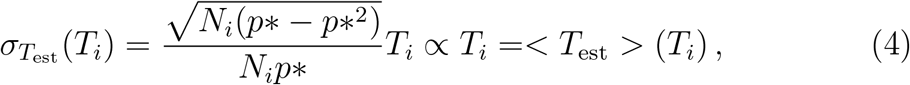

which is the statement of the scalar property. The proportionality constant, the Weber fraction, can be approximated by 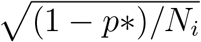 for *p* close to one.

To verify that the binomial distribution provides a good approximation to our data, Fig. 13 shows the relation between the mean and standard deviation of the firing probabilities *p*_*i*_ of each pool. The dotted curve represents the relation predicted by the binomial distribution. The probability *p*_*i*_ is estimated by normalizing the mean firing rates *FR*_*i*_ to the maximal value in each pool. The standard deviation is fitted to the relation from the binomial distribution, 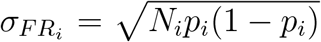 and the values in the figure are divided by the fit parameter 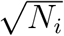. The fit is best for pools representing short intervals. For intervals above 400 ms, the firing probability *p*_*i*_ decreases and thus, the standard deviation increases. The estimated *N*_*i*_ values from the fit match the actual pool sizes well (estimate from the fit to the binomial distribution: 16 ± 5, average actual pool size: 14 ± 5) with no statistical difference (two-sample t-test: *t* (28) = 0.936, *p* = 0.36).

**Figure 13:**
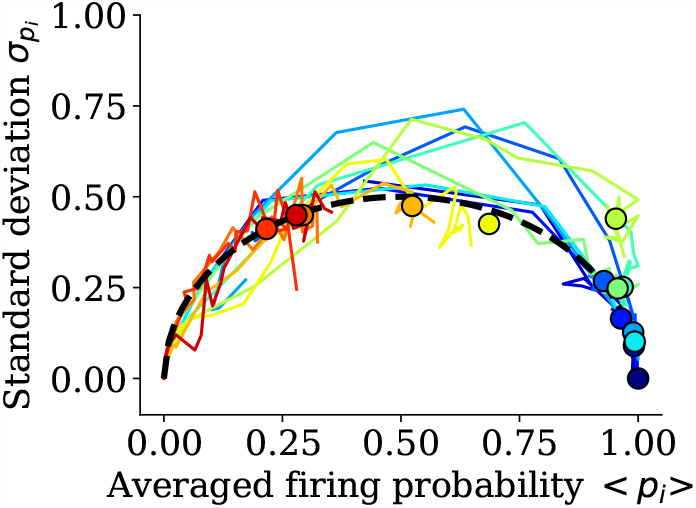
Binomial distribution of firing rate probabilities. Standard deviation of the firing probabilities as a function of the mean. The curves are well-fitted by the relation predicted by the binomial distribution (dotted line). For longer intervals, smaller firing rates and higher standard deviations occur. The colors correspond to the intervals that are represented in each pool (see the color bar, cf. Fig. 2A).

In the supplementary materials, see SAppendix B, we illustrate how the above computations can be generalized to arbitrary durations *t* and the full range of neurons. Importantly, this generalization requires that the shape of the output tuning curves *O*_*i*_(*t*) are largely conserved across output neurons *i*. In the following section, we discuss consequences of violating this assumption, which apparently occurs at longer durations (cf. Fig. 2A). Finally, we discuss how the stereotypic firing rate profiles (Fig. 8C) for the different IEPs of the network may arise.

The profiles approximate the cumulative normal distribution, which follows from the fact that each neuron fires as soon as its particular firing threshold is crossed, which is governed by a stochastic process: Given that the membrane potential of each neuron is driven by noise that can be approximated by a normal distribution, the probability to cross the threshold at a given time is the cumulative normal distribution with the difference of the mean membrane potential *< V*_*i*_(*t*) *>* from the threshold *V*_th,*i*_ as the mean. Note that this difference is decreased by the second stimulus, and we assume that the inputs are balanced such that the membrane potential is well below the threshold without the second stimulus, but close to the threshold in the presence of the second stimulus. In the following, we only consider the membrane potential under the influence of the second stimulus and assess under which conditions it may cross the threshold.

The standard deviation of the membrane potential does not systematically change over time, as neurons are driven back toward their resting potential by leak currents, which can be described by the Ornstein-Uhlenbeck process [82] for which the standard deviation is constant in time. Thus, the change in *p*_*i*_(*t*) over time is mainly driven by changes in the mean membrane potential *< V*_*i*_(*t*) *>*. The main driver of this change over time is the slow GABA_B_ inhibition after the first stimulus, which effectively prevents neurons from firing in response to a second stimulus within a certain window of time. Apart from the time constant of the GABA_B_ conductance, this time window is determined by the neuronal and synaptic parameters within each pool. As we have seen in Fig. 10, the mean current difference to the neurons’ rheobase increases for pools encoding longer intervals (which can be translated into the difference between the average membrane potential and the firing threshold). Thus, more of the inhibition from the first stimulus must be worn off before those pools can respond to the second stimulus, which happens at later times. In summary, the increase of *p*_*i*_ over time is mainly due to a gradual decay of long-term inhibition which decreases the difference between average membrane potential and firing threshold and thus, increases the chance of each neuron to fire. The different parameters in each pool determine the speed by which *p*_*i*_ increases over time.

To verify that the observed scaling of *p*_*i*_ can be reproduced by the mechanism described above, we have constructed a minimal model simulating a single neuron that receives a large amount of random, balanced excitatory and inhibitory currents and is subject to a GABA_B_ current following the time of the first stimulus and to a fixed depolarization at the time of the second stimulus (see Methods for details). Simulating this neuron for a large number of trials allows estimating *p*_*i*_ for a given firing threshold *V*_th,*i*_. For simplicity, we simulate the different pools by varying *V*_th,*i*_, although the above results suggest a variation of the synaptic properties. For each simulated pool *i*, we compute the first time *T*_*i*_ at with *p*_*i*_ exceeds 95 % and record *T*_*i*_ as the represented duration of this pool. When time is scaled by this duration, *p*_*i*_(*t/T*_*i*_) shows the same cumulative normal distribution time course for each pool (Fig. 14), as we see in the network model (Fig. 8D).

**Figure 14:**
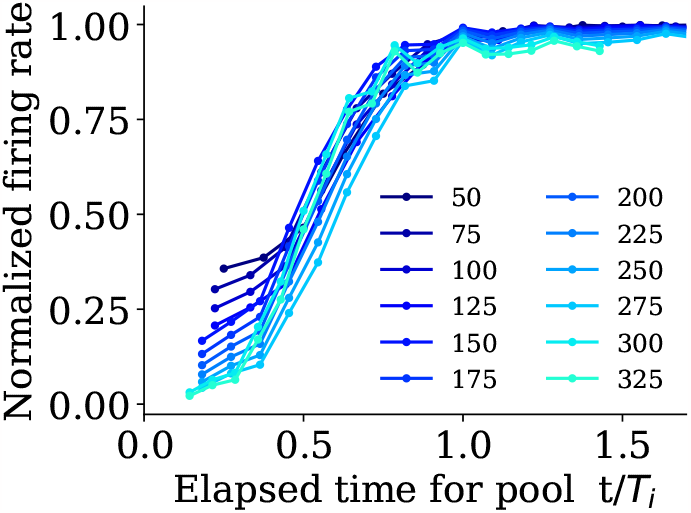
Results from the minimal model. Normalized firing rate as a function of simulated time, shown as a fraction of the encoded interval duration of each pool (color-coded). Each curve was generated by a different threshold for the membrane potential to elicit a spike. Higher thresholds lead to longer durations that need to elapse before that first spike (cf. Fig. 10). When scaling this time by the duration *T*_*i*_ at which the neuron spikes with 95 %, the firing rate curves for all durations largely overlap, as in Fig. 8D for the full model.

### Deviations from the scalar property and the origin on Vierordt’s law

As mentioned in the previous section, the scalar property relies on the assumption that the output tuning curves have the same shape for all output neurons. From Fig. 2A, it is apparent that this assumption is violated for output neurons representing intervals that are longer than 350 ms, where the tuning curves quickly grow broader as the interval durations increase. The widths of these tuning curves reflect the standard deviation of the underlying firing rate curves. Indeed, Fig. 13 shows that for longer intervals, higher standard deviations of the firing rates occur as the mean firing rates decrease and thus, the standard deviations move towards the middle of the half-circle implied by the binomial distribution. Here, we discuss the consequences of violating the assumption of stereotypic tuning curves of the output neurons.

If we consider the most extreme case where all curves *O*_*j*_ are completely flat such that all intervals contribute to the estimate with the same weight, it follows from the normalization of *O*_*j*_(*t*) to one that for each time *t*^∗^, the same duration estimate *< T*_est_ *>* is generated independently of the real time *t*^∗^, which is the average over all represented intervals *T*_*j*_. As the standard deviations are linked to the mean duration estimate, the standard deviations are also constant in time in this case. In intermediate cases, combining constant, sharp tuning curves *O*_*j*_ for short intervals *T*_*j*_ and increasingly broader curves for longer intervals, the scaling of both the mean and standard deviation are shifted away from the linear relations described in Eq. B.1 and B.4 towards constant values. This implies that deviations from the scalar property and Vierordt’s law, the observation that long intervals are underestimated and short ones are overestimated, are mechanistically linked. Indeed, for most manipulations described above, a strong negative linear relation is found between the slope of Vierordt’s law and the minimal standard deviation of the output curves *O*_*j*_, see Fig. 15 (*r* = −0.96, *t*(12) = −11.88, *p <* 0.01).

**Figure 15:**
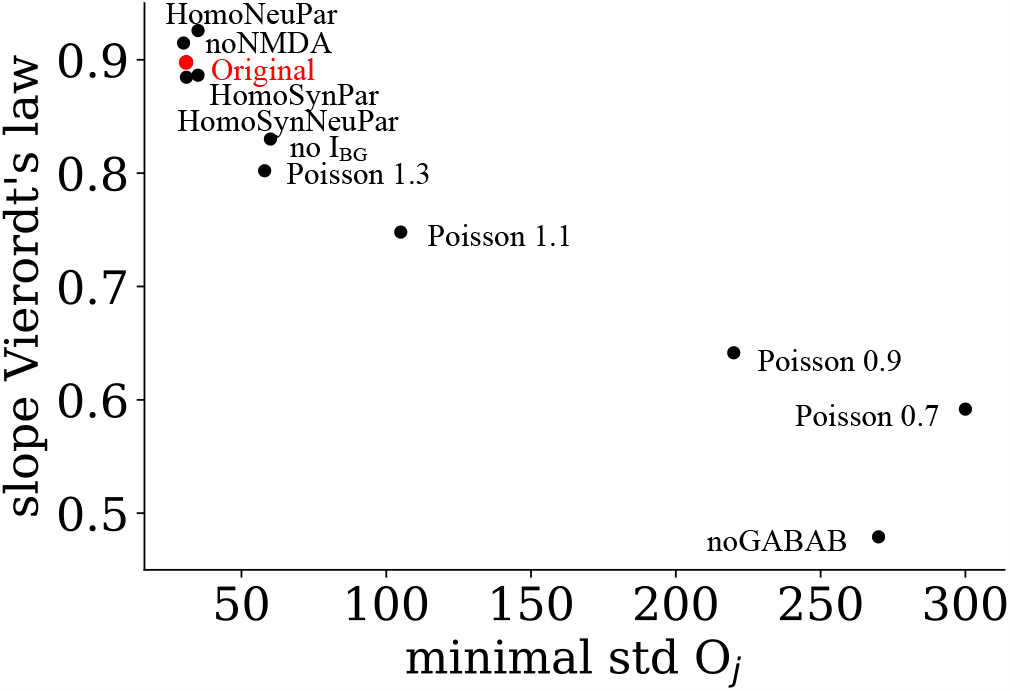
Relation between Vierordt’s law and the scalar property. Comparing the minimal standard deviation of the output curves *O*_*j*_ against the slope of Vierordt’s law for the different cases, results in a strong negative linear trend. The smaller the standard deviation, the better the estimation.

**Figure 16:**
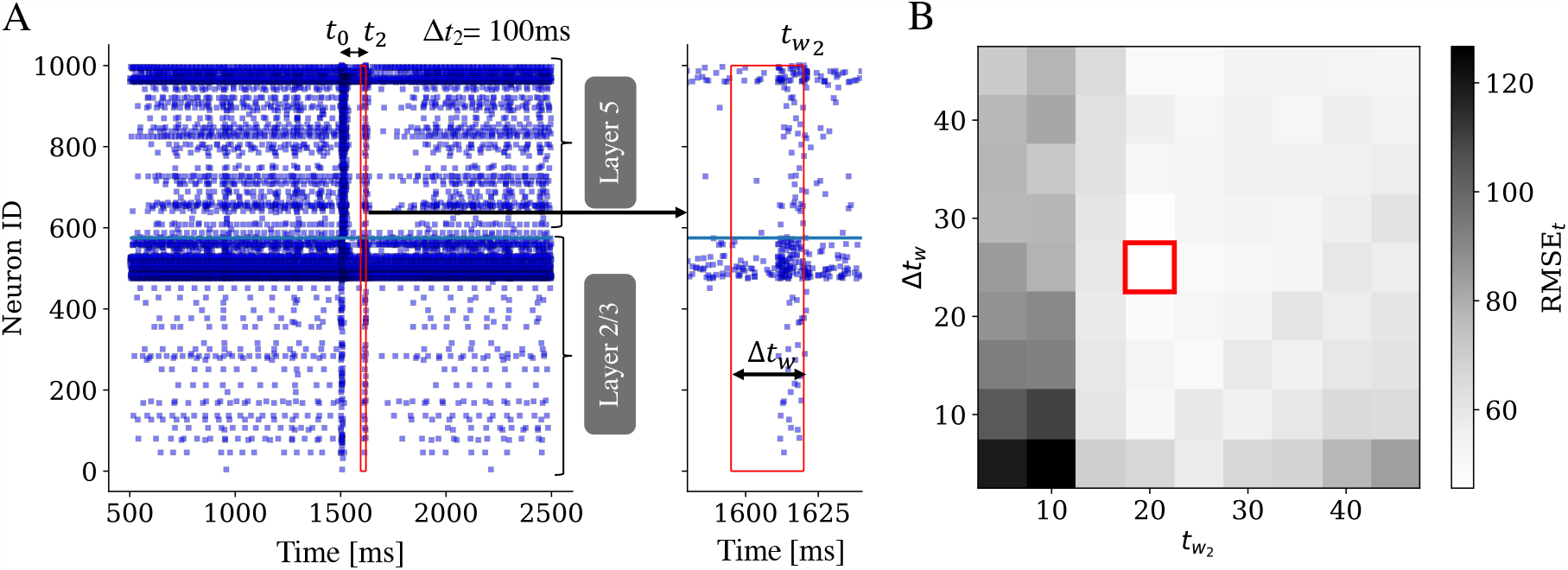
Extracting the states by defining a window. **A**. Rasterplot of all 1000 neurons with two stimulations (*t*_0_ = 0 ms for 10 ms and at *t*_2_ = 100 ms for 10 ms) to reproduce an interval of ∆*t*_2_ = 100 ms. The red box identifies the window in which the states from the spike trains are extracted (right: zoomed-in view). **B**. Result of a grid search to find the optimal parameters for the window with varying window size and end time. The best parameter set was selected by the minimum RMSE value (red box).

## 3. Discussion

We investigated whether the established state-dependent network model of subsecond timing [15] still works when embedded in a data-driven network model of the prefrontal cortex [27] and whether it is capable of reproducing some of the most important experimental results on time perception, namely the linear psychophysical law [35], the scalar property or Webers law [40] and the modulation of subjective duration by DA [2]. This investigation can be seen as an experimental validation of the model, which has not been conducted so far for any theory of time perception. Indeed, we showed that the model can successfully reproduce the linear psychophysical law and the Webers law for intervals in the range of 50 - 750 ms. Modulation of D2 DA receptor was tested by modifying neuronal and synaptic parameters and yielded, within a smaller range of 200 - 600 ms, a slowing of the internal clock for an antagonistic modulation (up to − 50 %) and, conversely, a speeding of the internal clock for an agonistic modulation (up to 20 %). As we discuss below in more detail, similar limitations are also observed in experiments. Overall, the model can be seen as validated by all of the chosen comparisons with experimental data. While those findings are among the most established in the literature on duration estimates, it would be desirable to test the model against further experimental evidence to further strengthen its position as a plausible mechanism for time perception in the brain.

In the following, we discuss the dynamic mechanism underlying subsecond timing in the state-dependent model, as well as the implications of the present work for explaining the three above-mentioned experimental results. Finally, we compare the current model to the original state-dependent network model by Buonomano [15].

### Mechanism and limitations of subsecond timing within the state-dependent PFC model

We found that different interval durations are encoded in specific populations of neurons (interval-encoding pools, IEPs) that were identified by the readout neuron onto which each neuron projects with the largest readout weight. Each duration-specific neuron pool shows a stereotypic firing rate response to the second stimulus that gradually increases as the interval becomes longer, up to a maximum just before the interval duration the pool encoded. These curves are reminiscent of the psychometric response curves that are seen in many time perception paradigms [83]. Importantly, the curves show very similar profiles when the time elapsed is scaled to the duration each pool represents (Fig. 8C). This similarity allows the readout neurons to decode the elapsed time of the firing rate of the neurons (the “states”) of the network at a given time. The scale invariance in the response of the pools is also an important prerequisite for the scalar property of timing errors, as we discuss in more detail below.

As we illustrate using a minimal model, the stereotypic response can be reproduced by a single neuron that is subject to random synaptic input and slow inhibition. For a given mean and standard deviation of the input, the firing of the neuron is governed by the binomial distribution. If the mean synaptic input increases over time, the firing rate approximates a cumulative normal distribution (Fig. 14). The scaling of this distribution is determined by the distance of the mean input current from the neuron’s rheobase, relative to its standard deviation. Thus, two ingredients are needed to explain the shape of the firing rate curves (and thus, the ability of the model to encode durations within a certain range): A synaptic current that changes systematically over time and a substantial variety of neuronal and synaptic parameters to allow different neurons (or pools of neurons) to encode different interval durations.

We conducted a series of correlation and ablation analyses in the full network model to identify the sources of these two factors. First, we found that the distance of the average current from the rheobase (Fig. 10B) increases for pools that encode longer intervals, consistent with the theory derived from the minimal model. Regarding the source of the systematic change in the input current, ablation analyses identified the slow GABA_B_ current as the most important component, as its ablation leads to a much more pronounced decrease of its slope in relation to a one-to-one relation (Vierordt’s law) and a higher trial-to-trial variability of duration estimates. The removal of shortterm plasticity (STP) leads to similar impairments but to a lesser degree (7A and D). Ablation of NMDA or removal of neuronal or synaptic heterogeneity did not have this effect (Suppl. Table A.3). On the other hand, removing GABA_B_ and STP also does not abolish the ability of the model to encode time altogether. So apparently, synaptic processes with shorter time scales (AMPA and GABA_A_) can also provide a time code, albeit with less accuracy (larger deviations between subjective and objective duration) and precision (larger standard deviations across trials).

We also assessed potential limitations of the model in terms of the length of the intervals that can be encoded and the levels of noise that can be tolerated. Already within the range of tested interval durations, we observed deviations both from the one-to-one psychophysical law (Fig. 2B) and the scalar property (Fig. 3) at interval durations above 500 ms. Both effects are even more pronounced when the network is trained for longer intervals, up to 2000 ms. Indeed, Fig. 6A and B suggest that there are two distinct linear regimes both for the psychophysical law and the scalar property for intervals above and below 500 ms, respectively. Finally, we observed a gradual broadening of the tuning curves of the readout neurons for intervals greater than 300 ms (Fig. 2A). Given the importance of GABA_B_ for precise and accurate subsecond timing, it is reasonable that these limitations are related to the time constants of GABA_B_. Indeed, the rise and decay time of that current add up to 300 ms, the interval duration where the tuning curves begin to degrade. Longer intervals could potentially rely on short-term synaptic plasticity, with time constants ranging up to 700 ms. For longer intervals, other timing mechanisms could play a role, since there is also experimental evidence to support that time estimation in the brain is performed by a number of different mechanisms in various brain areas for distinct interval durations [84, 85].

The gradual decrease in timing accuracy with increasing noise levels (Fig. 6C) as well as the deviation from the scalar property (Fig. 6D) can be explained similarly: As the levels of external noise increase, each output neuron becomes less predictive of the interval duration it represents, resulting in broader and shallower tuning curves, which are linked to both phenomena illustrated in Fig. 6C and D.

### Implications for experimental results on subsecond timing

*Linear psychophysical law*

The linear relation between subjective and objective time is among the most robust psychophysical findings in time perception [35] and thus, its fulfillment is seen as an important criterion for model validity. In the statedependent PFC model, the neural basis for the linear psychophysical law is the stereotypic firing rate profiles relative to the criterion time in each of the interval-sensitive pools of neurons. By extension, each of the readout neurons shows a similar tuning curve in response to the second stimulus, and thus, even stimuli that are presented at durations that are not explicitly trained for can be represented, as the (similar) tuning curves of the readout neurons encoding intervals nearby can provide a suitable interpolation. Small deviations from linearity can be seen at very short and very long durations (compare the dash-dotted line to data in Fig. 2B), resulting from boundary effects (incomplete tuning curves, see Fig. 2A, dark blue and dark red curves). Independent of these effects, a decrease in the slope of the linear psychophysical law can also be observed in the model, and thus, an overestimation of shorter intervals and an underestimation of longer intervals (Vierordts law [41, 42]). We showed that under the assumption of moderate, perfectly constant standard deviations of the tuning curves for all interval-selective output neurons, an exact one-to-one relation between subjective and objective time follows. In contrast, Vierordt’s law occurs if the standard deviations of the tuning curves increase for output neurons that encode increasing interval durations. This is the case for intervals exceeding the dominant time scale of the model, the combined rise and decay time of the GABA_B_ current, but also if this current is removed at all or increasing levels of external synaptic noise are added.

Previous studies have attempted to explain Vierordt’s law by finding that subjects overcome uncertainty during production tasks by optimizing their responses to the statistics of the previously shown trials. Fitting these results with various statistical models yielded the Bayesian framework as the best fit [86, 87, 88, 42]. A recent study analyzed the results from the classic experiments of Vierordt within such a framework [42] and found that much of the deviation from a one-to-one relation between subjective and objective time can be attributed to the specific experimental protocol used in these studies, namely a completely random variation of interval durations between trials. Within the Bayesian framework, a more naturalistic, gradual change of durations (using a random walk) abolished the distortion. Interestingly, when the authors performed a new experiment comparing both protocols, they found a residual distortion even in the random walk condition. The slope of the resulting psychophysical law (estimated in their Figure 3A) is approximately 0.86 [42], which is remarkably close to the slope we report for the current model without any adaptations (0.83, Table A.3). Thus, it seems likely that Vierordt’s law arises as a combination of Bayesian integration of information from previous trials and a distortion that is intrinsic to the network that generates the current duration estimate. As additional evidence for this proposal, note that the functional form of the psychophysical law (Fig. 2B) is similar to the one suggested by the Bayes least squares (BLS) model in [86] (their Fig. 3d), which also provided the best fit to psychophysical data.

Regarding the point of difference, the duration at which subjective and objective duration coincide despite a slope smaller than one, we found a relatively constant value around 500 ms for all different parameter sets as long as the same range of intervals (50 ms to 750 ms) is used to train the weights of the readout network. This value differs significantly from the arithmetic mean of all trained intervals (400 ms), which is supported by parts of the experimental literature [48, 49, 43], but not others [45, 43, 47]. However, the indifference point increases when a wider range of intervals has been used for training (50 ms to 2000 ms), suggesting that the indifference point can be shifted by training.

### The scalar property

The scalar property (or Weber’s law), the linear increase of the standard deviations of timing estimates with the mean duration, is an important experimental result that has often been cited as a cornerstone of time perception. Thus, any sensible model of time perception should be able to reproduce the scalar property, at least for a limited range of durations [89].

For the present model, we expected a sublinear instead of linear scaling of timing errors, since such results were reported in a study on one of its variants [26]. Surprisingly, the scalar property did emerge in our version of the model (Fig. 3), up to durations that correspond to the time constants of the GABA_B_ current. The importance of this current for the scalar property is underlined by the fact that its ablation leads to a drastic decrease of the cut-off duration, where Weber’s law is replaced by a constant timing error (Table A.4). We also found a plausible explanation for the origin of Weber’s law, combining profiles of firing probability that are scaled by the duration represented in each time-specific pool of neurons with a stereotyped temporal receptive field occurring from optimizing the readout from each of the pools in downstream units, both resulting in largely uniform shapes of temporal tuning curves in these output neurons and finally constant Weber fractions. In this regard, the mechanism generating Weber’s law bears similarity to the scalar filter banks in Grossberg’s spectral timing theory [90] and also to the multiple synfire chains in [17], each of which accounting for a given time interval and contributing to a timing error that scales approximately linearly within a certain range of durations. In contrast to these studies, however, the pools of neurons that represent specific durations spontaneously arise from the heterogeneity in the synaptic parameters, rather than being imprinted into the model architecture.

Several previous studies have attempted to explain the emergence of Weber’s law from general mathematical principles. Hass and Herrmann [57] have developed an information-theoretical framework to characterize the ability of a given model to reproduce the scalar property. They suggest that the sublinear (instead of linear) scaling of timing errors seen in many models may be caused by a common set of timing mechanisms that are based on the systematic change of some observable quantity in time, such as the firing rate of a neuron or spike in a specific neuron at a given time. The scalar property, on the other hand, is thought to emerge in models that use the increase of standard deviations itself (in a diffusion sense) to estimate time. The latter category comprises models that rely on purely stochastic transitions between neural states [91, 92, 93]. For example, Simen et al. [93] describe the slow ramping of firing rates by a specific form of drift-diffusion process that is driven by state transitions following a Poisson distribution. The scalar property then follows from the fact that the time between two Poisson-distributed events follows an exponential distribution, which has a constant coefficient of variation (equivalent to the Weber fraction) of one. In the state-dependent PFC model, timing is not driven by purely stochastic forces, but by a systematic change in excitability that is mainly driven by the time constant of the GABA_B_ current. Thus, it cannot be classified as a variance-driven model in terms of Hass and Herrmann [57]. Rather, the scalar property is generated by the stereotypic firing rate curves and the fact that the mean and standard deviation of timing estimates always covary in a specific way, governed by the binomial distribution. One could characterize the state-dependent model as a covert mean-driven model, as the systematic change in excitability is not directly observable.

Note that timing errors saturate into a constant of longer intervals, deviating from Weber’s law (Fig. 3). This kind of deviation is only rarely observed in experiments [52], while a more common observation is a steeper, superlinear increase in the error over time [52, 89]. A possible explanation for this discrepancy may be the fact that we only examined a single mechanism of time perception at a time. In the brain, several of these mechanisms are likely to coexist, potentially each with its own optimal time scale [14]. For example, the mechanism of ramping activity [16] has been shown to represent durations of up to several seconds. If this mechanism coexists with the state-dependent mechanism we investigated here, it is likely that the overall time estimate is taken over by ramping activity at intervals that are too long to be represented by the state-dependent network. Under this assumption, the constant timing error predicted for longer durations is not seen because, for those intervals, another, more suitable mechanism takes over. Theoretical frameworks for the integration of different sources of time estimates have been previously proposed [94], including our own proposal of a “temporal hub”, potentially located within the basal ganglia [14, 95]. Assessing how the combination of timing mechanisms with different time scales in such a hub would affect the scaling of timing errors seems to be an important direction for future research.

Finally, we note that the deviation from Weber’s law and the deviation of the slope of the psychophysical law from one (Vierordt’s law) are based on the same mechanism, at least in this model. Regarding experimental studies, this connection suggests that it may be worthwhile to study both laws together when searching for transitions between different mechanisms of timing or ablating specific neuronal of synaptic mechanisms in real neural systems.

### Dopaminergic modulation

The third major experimental result we aimed to reproduce in this study was the modulation of subjective duration by dopamine (DA). We modeled the effects of the D2 dopamine receptor on synaptic and neuronal properties based on results from *in vitro* experiments and, indeed, found that subjective duration is lengthened by D2 activation and shortened by D2 inactivation (Fig. 4), as it has been consistently seen in experiments [2, 62]. As shown in Fig. 4B, the change in subjective duration is mostly driven by the modulation of the synaptic parameters, namely a 20 % decrease in NMDA and 50 % decrease in GABA peak conductances at 100 % D2 activation compared to 0 % (see Methods for details). As GABA decreases much more compared to NMDA, activation of D2 results in a net increase in network excitability. As time is encoded by means of the firing rates in each of the pools in response to the second stimulus (Fig. 8D), increased excitability implies that any given firing rate will be reached faster under D2 activation and thus the subjective duration will be shorter than the objective duration. The opposite is true for D2 inactivation.

Understanding this mechanism of dopaminergic modulation of subsecond timing also helps to explain how the model reproduces a number of effects that are also seen in experiments. First, the scalar property is unaffected by mild D2 modulation (Fig. 4C), consistent with experimental results [96]. As D2 activation (or inactivation) simply changes the excitability of the entire network, neurons in all pools are shifted towards earlier (or later) responses, while the relation between means and standard deviations of firing rates that cause the scalar property remains intact. Second, the retraining of the output neurons fully compensates for the modulation of D2 (Fig. 5). This is consistent with experimental findings on the chronic application of dopaminergic drugs such as methamphetamine to rats [97] and also, more broadly, results suggesting neuroadaptive processes in response to chronic activation of the dopaminergic system, e.g., by addiction [98, 99]. We can explain this effect by a synaptic mapping of neurons within the network that changed their optimal reaction to the second stimulus to an earlier or later time under chronic dopaminergic influence. For example, if neurons in a pool that encode 100 ms are sensitive to 150 ms under D2 inactivation, retraining would assign them to the output neuron that encodes 150 ms. Thus, the change in subjective duration can be fully compensated if the modulation is present long enough such that the synaptic connections to the output neurons can rewire accordingly. This is the case if the activation or deactivation of D2 is chronic and stable over time. Finally, we observed limitations in both the durations of the intervals and the activation and deactivation levels of D2 within which the subsecond timing works properly. In particular, for D2 activations beyond 50 %, as well as interval durations shorter than 200 ms, the subsecond timing basically stops working. In Suppl. Fig. A.4, we see that all time estimates collapse to a single value for values far beyond these boundaries. Deactivation of D2 shows similar effects, but subjective durations can still be differentiated for intervals above 200 ms at deactivations of − 100 % (Suppl. Fig. A.4). Indeed, experimental results using D2 receptor agonists point in the same direction as our findings on D2 receptor activations [96]. More specifically, the study shows that the scalar property breaks down at higher concentrations of the agonist. Within our model, we can explain these results as a boundary effect: An increase in D2 activity makes the entire model overly excitable. Thus, the different pools lose their ability to differentiate between different intervals and the timing over a range of intervals is impaired. This proposal is supported by the fact that subjective durations for the different pools collapse to a single value as D2 activation approaches 100 % (Suppl. Fig. A.4) and that subjective durations in pools encoding short intervals (dark blue lines) do so for smaller D2 activation levels, as they are more excitable to begin with.

It is important to mention that, in contrast to the agonistic modulation, the antagonistic DA modulation could only be extrapolated by inverting the parameter changes of the agonistic modulation, due to the lack of experimental data. This inversion can be interpreted as a deactivation of tonically active D2 receptors. Although it is not guaranteed that such a deactivation will yield the exact opposite of the activation, it seems unlikely that the net effects on synaptic conductances will be qualitatively different, given the large differences of the effects on NMDA and GABA for the agonist. Nevertheless, future experiments should explicitly measure the parameter changes for DA antagonists to validate our assumptions.

Although there is sufficient evidence for a D2 receptor-based dopaminergic modulation of duration estimates in the subsecond range [66, 67, 64, 68, 69], most studies have been performed in the seconds to minutes range [2, 65]. However, we emphasize that the basic mechanism of dopaminergic modulation is not necessarily restricted to the subsecond range. As D2 activation results in a net increase in network excitability, any duration estimate based on neuronal or synaptic activity will be affected in the same way as described above. In particular, we could think of additional neuronal or synaptic processes with longer time scales that extend the range of interval durations the state-dependent network model could represent.

Finally, note that we have restricted our analysis to the D2 dopamine receptor, as most effects on subjective duration have been reported in relation to this receptor type [2] and it has been shown that the binding affinity of a variety of drugs to the D2 receptor predicts the effect on time perception [63]. Moreover, the effects of D1 receptor activation are likely more complex than a simple change in excitability [100, 101], so we would not expect a systematic change in subjective duration within this model. However, it should be noted that more recent studies also showed effects of D1 on time perception, in particular in the context of ramping activity [102]. Therefore, it would be worthwhile to investigate the underlying mechanisms of this modulation as well and to assess the circumstances under which the modulation of D1 and D2 is the most important, respectively.

### Relation to the original state-dependent model

Here, we compare our state-dependent PFC model with the original statedependent model [15]. Using 500 neurons, [15] showed normalized output spikes up to 250 ms when 50 ms steps are used for training and spikes up to 400 ms when 100 ms steps are used. Our time constants for GABA_B_ are similar, as they have the same rise time *τ*_*rise*_ = 100 ms, but a slightly longer decay time *τ*_*decay*_ = 200 ms based on the electrophysiological literature [76] (compared to *τ*_*decay*_ = 166 ms in [15]). Within the range 50 - 250 ms, we were also able to show similar output patterns, but for longer intervals, our output peaks become broader. However, our model was able to perform discrimination tasks up to 750 ms.

For the original state-dependent model, Buonomano showed reduced timing performance when GABA_B_ or STP were removed, consistent with our results. Furthermore, we showed that reducing the heterogeneity of STP by allowing only one type of STP within each given neuron type combination greatly impaired timing performance. To understand the phenomenon, note that the dominant STP type for E-to-E connections is facilitating, while for I-to-I connections, it is depressing. Thus, if we limit STP to a single, preferred type for each connection types, the network is put into a regime where excitation is facilitated and inhibition is depressed at the same time. This combination will likely make the network excessively excitable as a whole, affecting the necessary differentiation in excitability between pools. In a recent study of persistent activity within the same PFC network [103], we observed that persistent activity stabilized when the STP was homogeneous compared to no STP at all (and also heterogeneous STP, which prevented persistent activity at all). This finding also supports the idea that the excitability of the network is enhanced by homogeneous STP, similar to excessively strong activation of the D2 receptor (see previous section).

Furthermore, Buonomano claimed that temporal tuning was specifically achieved by the variability of synaptic weights, and homogenous synaptic weights prevented interval discrimination. On the contrary, when we performed simulations with homogeneous neuronal and synaptic parameters, timing could still be achieved. Apart from synaptic weights, connectivity might also play an important role in our timing tasks. For instance, neurons that receive few negative inputs from inhibitory neurons can easily be reactivated for a second interval compared to neurons that receive more inhibitory inputs.

In summary, timing of a range of different intervals requires a process with long time constants such as GABA_B_, and heterogeneity in the parameters that determine the excitability of a neuron. Even if one component is removed, there might be other heterogeneous parameters within the network that help to differentiate the neurons to obtain interval discrimination, given sufficient heterogeneity. Furthermore, we note that GABA_B_ is not the only option for enabling subsecond timing in the range of hundreds of milliseconds. In principle, any process with a sufficiently long time constant could be used, for example, the influx of calcium ions following a spike that has already been explored in the context of the ramping activity model [16].

The fact that the state-dependent PFC model showed the scalar property was surprising, since a previous study on the original state-dependent network model showed a sublinear rather than a linear scaling of timing errors [26]. We showed that timing errors (averaged over several combinations of parameters) are best fitted with a linear part for shorter intervals and a constant part for longer ones (Fig. 3). For any individual parameter combination, the constant part at longer intervals is less clearly visible (e.g., Fig. 2C) and may be mistaken for a decreasing slope in an overall sublinear scaling. Taken together, these results suggest that a reanalysis of the previous data, pooling different parameter sets and considering a saturation of timing errors at longer intervals, may yield results that are similar to ours. On the other hand, the intervals considered in [26] regarding timing error did not exceed 200 ms, a duration in which a sublinear increase of the standard deviation may be due to the generalized form of Weber’s law [53, 104]. On the contrary, the intervals considered in the present study are sufficiently long to examine the scalar property.

### Conclusion

Computational models provide opportunities to understand the neural mechanisms that underlie time perception. We chose a set of key experimental findings to test such models against, namely, the psychophysical law (including Vierordt’s law), the scalar property of timing errors, and the dopaminergic modulation of subjective duration. The state-dependent PFC model is capable of reproducing each of these findings for interval durations between 50 and 750 ms, including the scalar property, one of the most enigmatic results in time perception. The model even reproduces several details of the experiments such as the magnitude of the Weber fraction and the slope of Vierordt’s law, the dependence of the indifference point upon the range of presented intervals, and the effect of the D2 dopamine receptor on the scalar property. Remarkably, no explicit tuning of the network parameters was necessary to obtain these results, except for the synaptic weights for the read-out network. Beyond providing strong support for the present variant of the state-dependent model, these results also strengthen confidence in the underlying network model of the prefrontal cortex [27], which has already been shown to reproduce key features of *in vivo* neural activity without the need for parameter adjustment and produced unintuitive results with respect to working memory [34].

Taken together, the state-dependent model of time perception, originally proposed by Buonomano [15] and implemented here within a realistic PFC model, can now be considered the first timing theory validated by a range of important experimental results. However, this does not rule out alternative theories at all, especially for interval durations beyond 750 ms. Other prominent approaches such as models of ramping activity [16], synfire chains [17] and the striatal beat model [18, 19] can now be tested in the same way. This will likely lead to a range of validated theories of time perception and a more thorough understanding of how the brain tells time under different circumstances, such as different ranges of durations.

## 4. Methods

### Network model

To test timing within a realistic prefrontal cortex model, we used the spiking neural network model proposed by [27]. This model can reproduce key features of *in vivo* activity, as the neuronal parameters and synaptic parameters were constrained by anatomical and electrophysiological data from *in vitro* experiments on rodents. The spiking neural network consists of two laminar components, layer 2/3 and layer 5, of which 47.0 % within layer 2/3 and 38.0 % within layer 5 were excitatory pyramidal neurons and 10.4 % within layer 2/3 and the 4.6 % within layer 5 were interneurons. The interneurons comprise fast spiking cells, bitufted cells, Martinotti cells, and basket cells [27]. Neurons were modelled by the simplified adaptive exponential (simpAdEx) integrate-and-fire neuron model shown in Eq. 5 and 6 proposed by Hertaeg and colleagues [73].

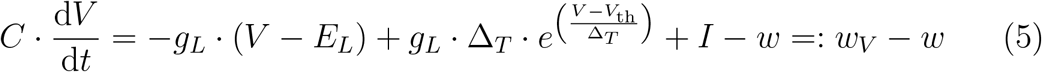

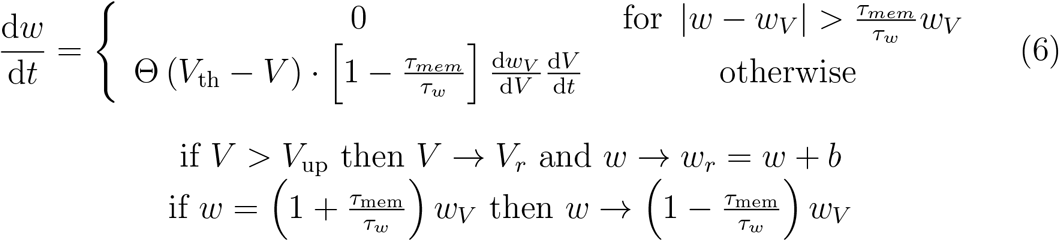

C describes the membrane capacitance, *g*_*L*_ the leak conductance, *E*_*L*_ the reversal potential, and *τ*_mem_ and *τ*_*w*_ the membrane and adaptation time constants. The differential equation for the membrane potential *V* includes an exponential term with a slope parameter ∆*T* . This term causes a strong increase of *V* once the membrane threshold *V*_th_ is passed [73]. *w* represents the spike-triggered adaptation variable with an additional differential equation with Θ being the heavy-side function and *w*_*V*_ the V-nullcline of the system (Eq. 5). If *V* crosses the peak potential *V*_*up*_, the membrane potential is set to the reset potential *V*_*r*_ and *w* is increased by *b* [73].

The network consists of 1000 simpAdEx neurons with parameters randomly drawn from experimentally validated distributions and pairwise connection probabilities from the literature [27]. Synaptic connections were simulated by conductance-based synapses for different receptor types *X* ∈ *{*AMPA, GABA_A_, NMDA}, where the postsynaptic current *I*_syn_ of a presynaptic spike train *{t*_sp_*}* is calculated using double exponential functions and including the reversal potentials for each receptor type 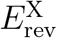:

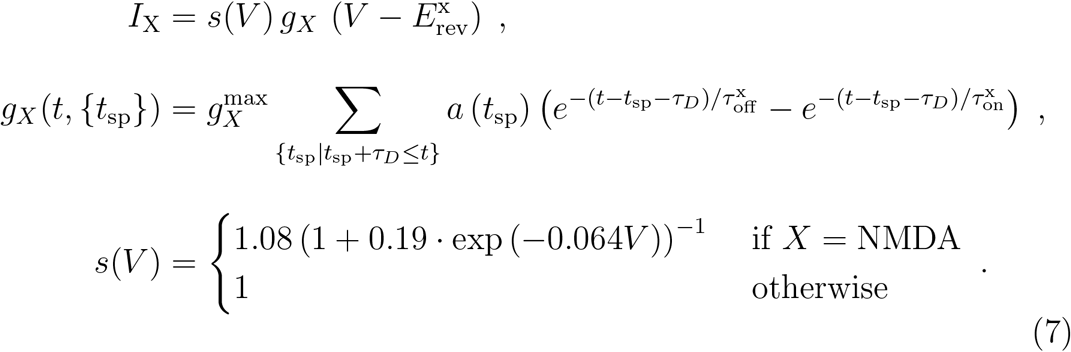

*g*_*X*_(*t, {t*_sp_*}*) describes the conductance dynamics with the second exponential describing the opening of the channel with a rise time constant *τ*_on_, while the first exponential accounts for the closing of the channel with a decay time constant *τ*_off_ . Both opening and closing are subject to a synaptic transmission delay *τ*_*D*_ and the NMDA synapses were simulated by multiplying the synaptic current by a nonlinear voltage-dependent term *s*(*V*) to simulate magnesium blockade at lower voltages, see Eq. 7.

Each synaptic connection is equipped with short-term plasticity via *a* in Eq. 7, implemented by the corrected version of [105]. Three different types of plasticity are implemented within the prefrontal cortex (PFC) model, namely short-term facilitation (E1/I1), short-term depression (E2/I2) and a combined version of both (E3/I3) [74, 75]. The cell types of the pre-and postsynaptic neurons determine which of these classes is used for each individual combination. In particular, connections among pyramidal cells, among interneurons, and between fast-spiking interneurons and pyramidal exhibit all three types of connections to varying degrees (pyramidal to pyramidal: 45% E1, 38% E2, 17% E3 [75]; interneuron to interneuron: 29% I1, 58% I2, 13% I3 [74], fast-spiking interneuron to pyramidal: 25% I1, 50% I2, 25% I3 [74]). A constant background current of *I*_exc_ = 250 pA was applied to excitatory neurons and a current of *I*_inh_ = 200 pA to inhibitory neurons. For more details, see [27].

The original model was supplemented with GABA_B_-type synapses to simulate slow IPSPs, which were found to be important within the state-space model [15]. The synaptic parameters for the conductance-based double exponential functions of GABA_B_ were extracted from Golomb et al. [76], see Table 4. The maximum conductance of GABA_A_ and GABA_B_ was also reduced by multiplying by a factor of 0.3 to counteract the excess inhibition within the network. The extended model still reproduces features of the original PFC model. In particular, the inter-spike intervals and the coefficient of variation (CV) per neuron are in the same range with and without GABA_B_ (see Suppl. Fig. A.1C, D. The time-averaged subthreshold membrane fluctuations of non-zero firing neurons are also consistent with GABA_B_ (*μ* = 3.1 mV, *σ* = 2.1 mV) and without GABA_B_ (*μ* = 2.1 mV, *σ* = 1.3 mV), cf. Suppl. Fig. A.1E.

**Table 4:** Synaptic parameters of GABA_B_.

| Synaptic Parameters | values |
| --- | --- |
| $E_{\text{rev}}$ | $-90$ mV |
| $\tau_{\text{on,GABA}_B}$ | $100$ ms |
| $\tau_{\text{off,GABA}_B}$ | $200$ ms |
| $g_{\text{peak,GABA}_B}$ | $\frac{1}{5} \cdot g_{\text{peak,GABA}_A} = 0.2$ nS |

To test timing within the PFC model, the full network as shown in Fig. 1 within the gray box was stimulated at the beginning and at the end of time intervals with a step current of *I*_s_ = 220 pA for 10 ms to test various inter-stimulus intervals 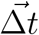. The model should ideally predict the presented inter-stimulus interval. All simulations were performed with the NEST Simulator (version: 2.18.0 [106]) after extracting the parameters from a routine in MATLAB R2020a [27].

### Training and testing the readout

After running the simulations, we extracted the states of all neurons (*N* = 1000) within a given time window ∆*t*_*w*_ following the readout mechanisms of a liquid state machine [77, 78] (cf. Fig. 1). Specifically, the states *s* in the PFC network were calculated using a sum over spikes within a small time window ∆*t*_*w*_ that ends 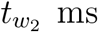 after the second stimulation (Fig. 16A) in which each spike is weighted using an exponential decay rate *τ* ^−1^, see Eq. 8.As *τ* gets smaller, the spikes which occur at the end of the window are more strongly weighted, and if *τ*→ ∞, the spikes of each neuron within the window are counted without any weighting.

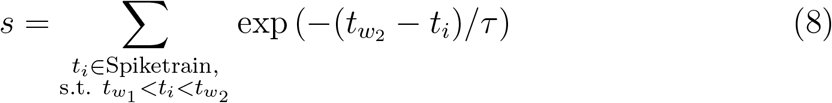

To predict interval durations, a readout layer was added to the model with the number of output units *N*_*r*_ matching the number of interval durations presented during training, cf. color-coded readout units in Fig. 1. Each output unit is then trained to be active for the respective time interval and to be inactive for all other time intervals. The readout weights were trained for all stimulus intervals and trials by maximizing the activity of the readout units for their corresponding stimulus intervals and minimizing activity for the remaining stimulus intervals.

The training of the readout weights was realized by calculating the least squares of the expected targets, cf. Fig. 1. After training with *N*_training,trials_ = 300 trials, the performance of time estimation was tested by presenting test intervals in 25 ms steps from 50 ms up to 750 ms and calculating the outputs of the previously trained readout weights. To predict the interval durations of the interstimulus intervals presented, the negative output values were first set to 0, then the positive values were normalized by the sum of all readout neurons, multiplied by the respective training time intervals, and summed across all readout units. The standard deviation of the estimated time was calculated on *N*_test,trials_ = 50 test trials. Including the negative output values in the estimate did not significantly change the results.

The hyperparameters that determine the states are the endpoint 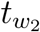 and the length of the window ∆*t*_*w*_ from which the spikes were extracted, and the exponential weight decay rate *τ* . These parameters were selected using a grid search and optimizing for minimal root mean squared error RMSE_t_ between subjective and objective time over a subset of training intervals (*N*_train, sub_ = 50) see Fig. 16B. In a first coarse grid search, we found *τ* → ∞, effectively summing up the spikes within the window, to give the lowest RMSE_t_ for all the settings of the other two parameters. Therefore, we fixed *τ* → ∞ and further optimized the two parameters ∆*t*_*w*_ and 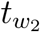in a finer scale grid search as shown in Fig. 16B. The optimal parameter sets were selected by the minimum root mean squared error ⟨RMSE_*t*_⟩_**trials**_ = 45.6 ms with ∆*t*_*w*_ = 25 ms, 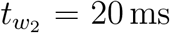and *τ* → ∞ as indicated by the red box in Figs. 16A, B.

### Dopaminergic modulation

The agonistic and antagonistic D2 receptor modulations were simulated by modifying various neuronal and synaptic properties. To alter neuronal parameters, we extracted the changes from our own unpublished data, in which the D2 receptor agonist, quinpirole, [107] was applied with a concentration of 10 *μ*M. On the contrary, changes in synaptic conductance were extracted from the literature (see Table 5). Specifically, we interpreted the changes in neuronal and synaptic modulation under the application of dopamine (DA) as 100 % agonistic modulation, cf. Table 5, the properties without application of DA as 0 % modulation and linearly interpolated between these two to simulate various DA concentrations in between.

**Table 5:** Parameter alterations for agonistic D2 modulation.

| Parameters |  | change | literature |
| --- | --- | --- | --- |
| **Neuronal parameters** |  |  |  |
| Reset potential | $V_r$ | -5.2% | |
| Threshold | $V_{th}$ | -3.7% | |
| Capacity | $C_m$ | +1.2% | |
| Leak conductance | $g_L$ | -14.2% | |
| Leak reversial potenital | $E_L$ | +6.6% | |
| Delta T | $sf$ | + 27.8% | |
| Peak Potential | $V_{up}$ | -4.1% | |
| adaptation time constant | $t_{cw}$ | -14.2% | |
| spike-triggeried adaptation | $b$ | + 9.8% | |
| **Synaptic parameters** |  |  |  |
| Peak conductance NMDA | $g_{NMDA}$ | -20% | [108, 109, 110] |
| Peak conductance GABA | $g_{GABA}$ | -50% | [111] |

Although experimental data for the change in neuronal parameters are available for DA agonists, the corresponding data for DA antagonists are lacking. Therefore, we assumed exactly opposite effects for DA antagonists and extrapolated the change in neuronal and synaptic parameters from 0 % to − 100 % to simulate antagonistic modulation via DA.

To measure the effects of an acute change in neuronal activity induced by DA, we applied the modulation at test time without retraining the readout weights, whereas for the examination of long-term plasticity-based adaptions induced by DA, we also retrained the readout weights in a second step for the case of ±30 % agonistic and antagonistic modulation.

### Noise

To simulate variability between trials, an input layer of Poisson neurons (*N*_Poisson_ = 10)) was added so that each PFC neuron on average received input from one Poisson neuron (*p*_poisson_ = 0.1) with a firing rate of *f* = 1 Hz. A typical cortical neuron receives connections from approximately 1000-10 000 other neurons [80, 81]. Therefore, we examined the effect of higher input variability while performing the limitation studies (see Limitations of the model in Results) in our model by increasing the firing rates of our Poisson process to *f* =1000 Hz, thereby emulating 1000 additional inputs. To test different noise levels, we increased the synaptic weights of Poisson neurons (*w*_Poisson_), and in parallel reduced the background current (*I*_back_) to prevent overexcitation (see Table 2 for details). For each synaptic weight factor, the background current was optimized using a grid search to minimize the RMSE of the average firing rates per neuron type and layer. The time-averaged subthreshold membrane potential fluctuation, i.e., after removal of spikes, was shown to be at 4 mV in experiments [112], which could already be reproduced within the PFC model [27]. For decreased background current and accordingly increased noise levels as described above, we find that subthreshold membrane potential fluctuations are within the range of 2 − 4 mV, cf. Suppl. Fig. A.7.

### Rheobase

The excitability of each PFC neuron was determined by rheobase *I*_rheo_ as proposed by Hertaeg et al [73] as shown in the equation. 9.

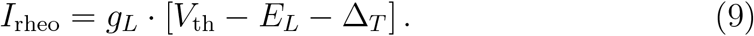

### Minimal model

In order to further understand how the network firing rate changes over time, we constructed a minimal model comprising a single neuron that is subject to the same synaptic currents and external inputs that are present in the model PFC. The neuron is modeled as a leaky integrate-and-fire neuron with parameters that are identical to the average values for an L23 pyramidal cell in the PFC model (*C*=170 pF, *g*_*L*_=5.85 nS, *E*_*L*_=−75.5 mV, *V*_th_=−65 mV). AMPA, GABA_A_, NMDA, and GABA_B_ synapses are modeled as in the PFC model (without STP). Except for GABA_B_, all synapses are stimulated by a Poisson spike train with *λ* = 0.25, simulating random spike trains in 4000 Hz for both excitatory and inhibitory synapses, balancing excitation and inhibition. To reproduce the effect of GABA_B_ after the first stimulation, the GABA_B_ synapse is only stimulated by 20 spikes during the first 20 ms of the simulation in response to the first stimulus that marks the beginning of the interval. This simplification is justified by the fact that the first stimulus elicits a much stronger effect on GABA_B_ compared to the ongoing activity and this strong initial activation is one of the essential features of the model that enables time perception. The neuron is subject to a constant background current of 40 pA, which is increased to 90 pA for 20 ms at the time of the second stimulus. The duration of the interval is varied between 50 ms and 500 ms in steps of 25 ms. We simulated *N* = 100 independent neurons and computed the mean and standard deviation of the membrane potential over all neurons at each of these durations. Assuming a normal distribution of the membrane potential, we then computed the probability *p* of the membrane potential to exceed the firing threshold *V*_th_ and interpreted *N*· *p* as the firing rate at a given time, which is depicted in Fig. 14. The different curves in this figure were generated by varying *V*_th_ between − 67 and − 63 mV in steps of 0.1 mV. Similar results are obtained by simply counting the number of neurons exceeding *V*_th_ in each case.

## Acknowledgments

The authors thank Daniel Durstewitz (Central Institute of Mental Health, Mannheim) for providing us with advice and computational resources. We also thank Jakob Jordan (Department of Physiology, University of Bern, Switzerland) for his help in implementing the neuron model in NEST, Tatiana Golovko (Central Institute of Mental Health, Mannheim) for collecting the dopaminergic modulation data, and Carolin Mößnang-Sedlmayr (SRH Hochschule Heidelberg) for helpful comments on the manuscript. The authors acknowledge the support by the state of Baden-Württemberg through bwHPC and the German Research Foundation (DFG) through grant no HA 7706/2-1.

## Appendix A. Supplementary materials S1

**SI Fig A.1:**
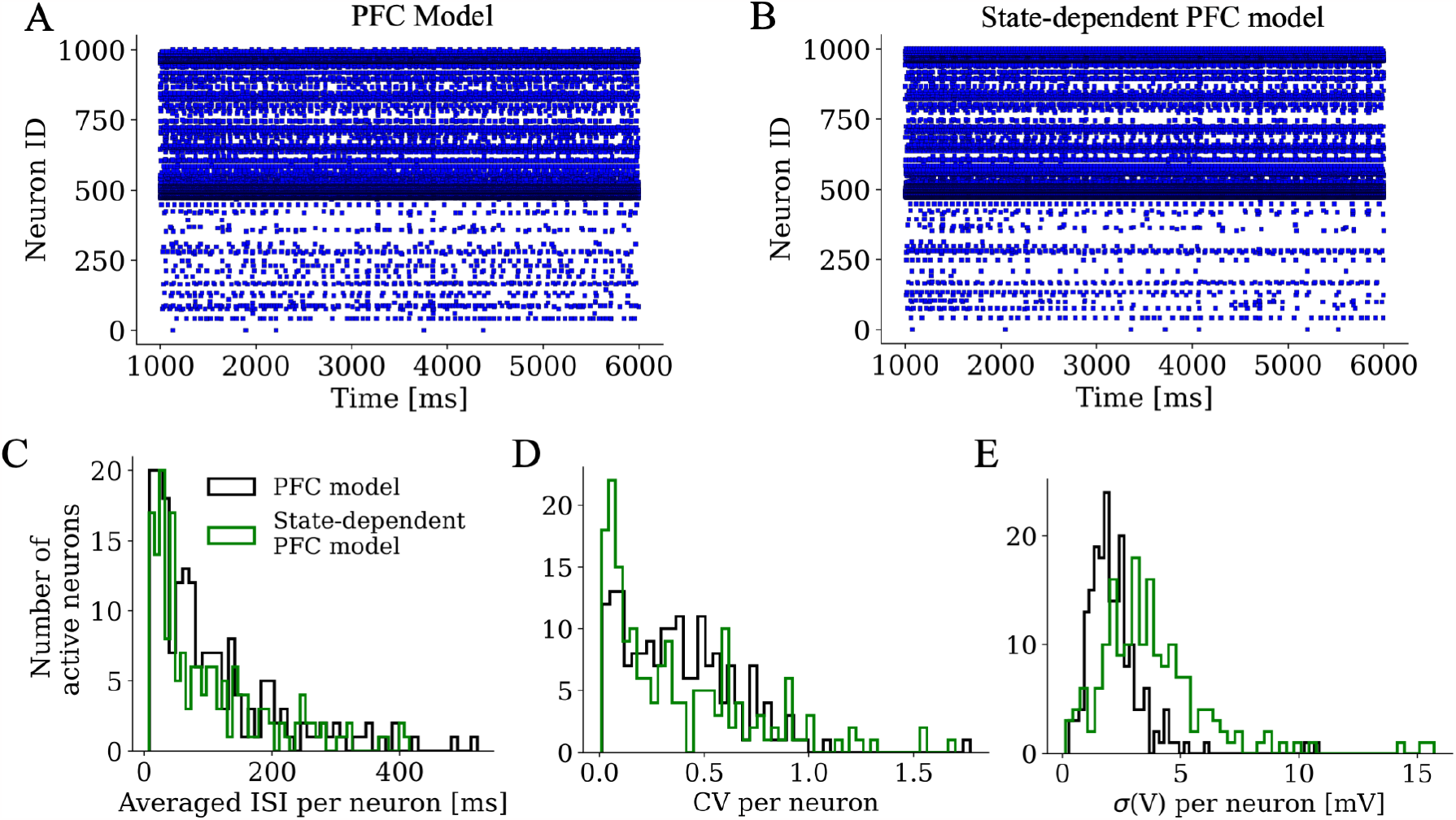
Matching spike statistics of the PFC and the state-dependent PFC model. The raster plots are shown for the PFC model without GABA_B_ in **A** and for the statedependent PFC model with GABA_B_ in **B. C** The numbers of active neurons are depicted as histograms for the averaged ISI for the state-dependent PFC model (green) (*μ* = 101.1 ms, *σ* = 93.2 ms and the PFC model (black) (*μ* = 106.3 ms, *σ* = 100.6 ms) for neurons with more than 10 spikes. **D** The CV per neuron for the same neurons (spikes *>* 10) is calculated for the PFC model in black (*μ* = 0.4, *σ* = 0.3) and for the state-dependent PFC model in green (*μ* = 0.4, *σ* = 0.4). **E** The standard deviation of the subthreshold membrane potential over time for non-zero firing neurons is higher for the state-dependent PFC (*μ* = 3.8 mV, *σ* = 2.4 mV) than for the PFC model (*μ* = 2.2 mV, *σ* = 1.3 mV) as shown with the histograms.

**SI Fig A.2:**
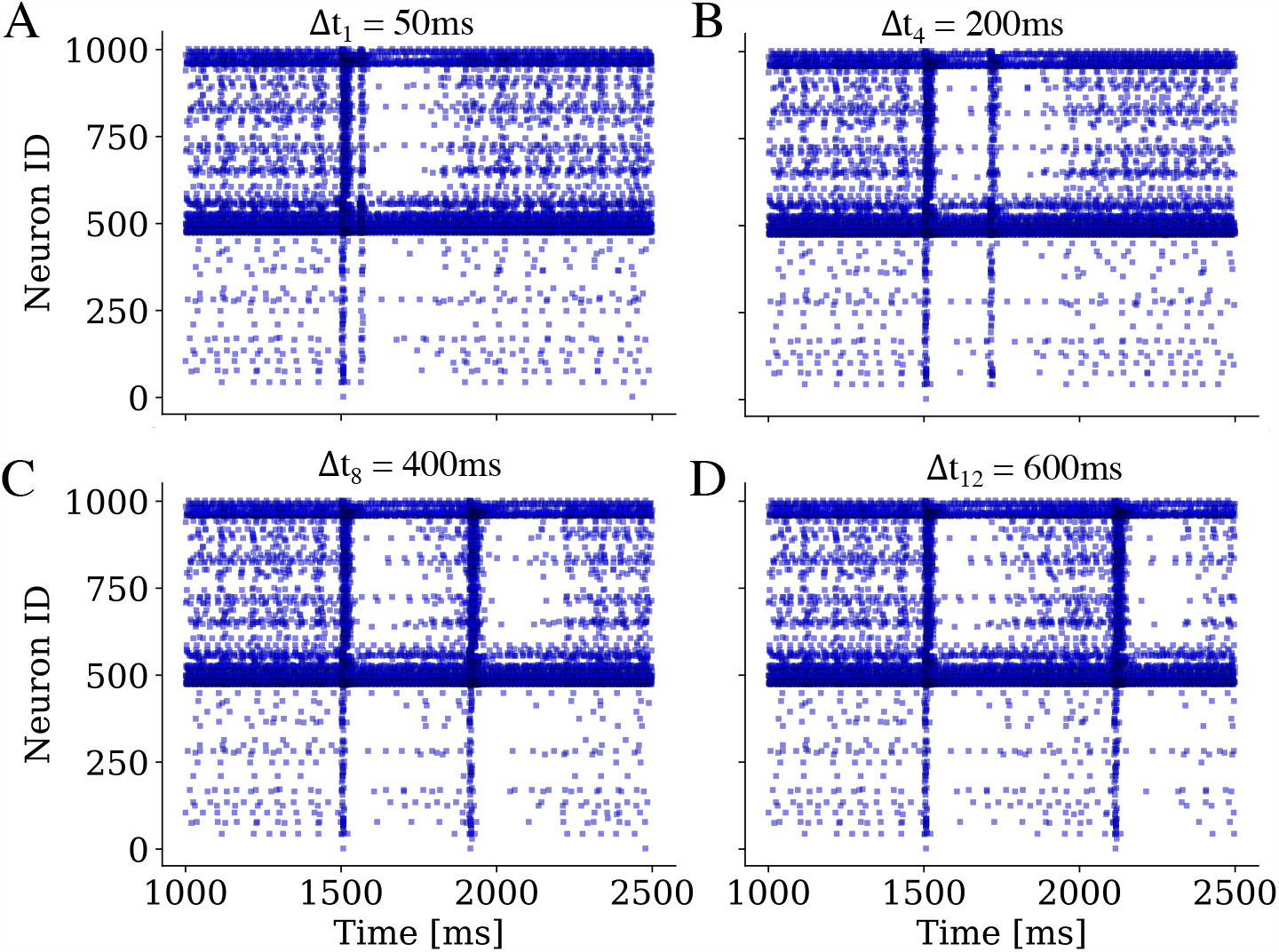
Spike trains for exemplary inter-stimulus intervals ∆t. The raster plots of the main simulation as shown in Fig. 2 are depicted for the training intervals ∆t_1_ = 50 ms in **A**, ∆t_4_ = 200 ms in **B**, ∆t_8_ = 400 ms in **C** and ∆t_12_ = 600 ms in **D**.

**SI Fig A.3:**
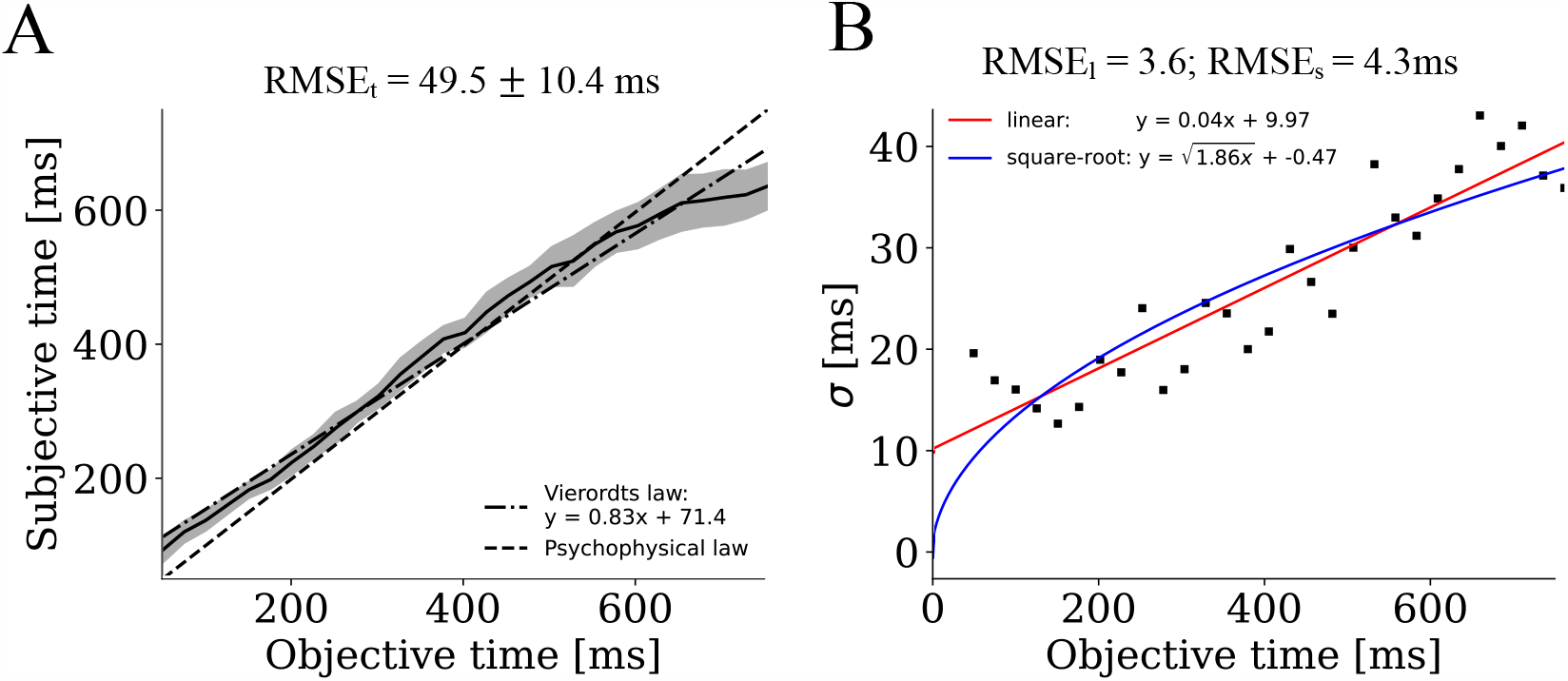
Ridge regression. Estimated times **A** and standard deviations **B**, when using ridge regression instead of the linear least squares method to calculate readout weights. Results are almost identical compared to the linear least squares method; cf. Fig. 2.

**SI Fig A.4:**
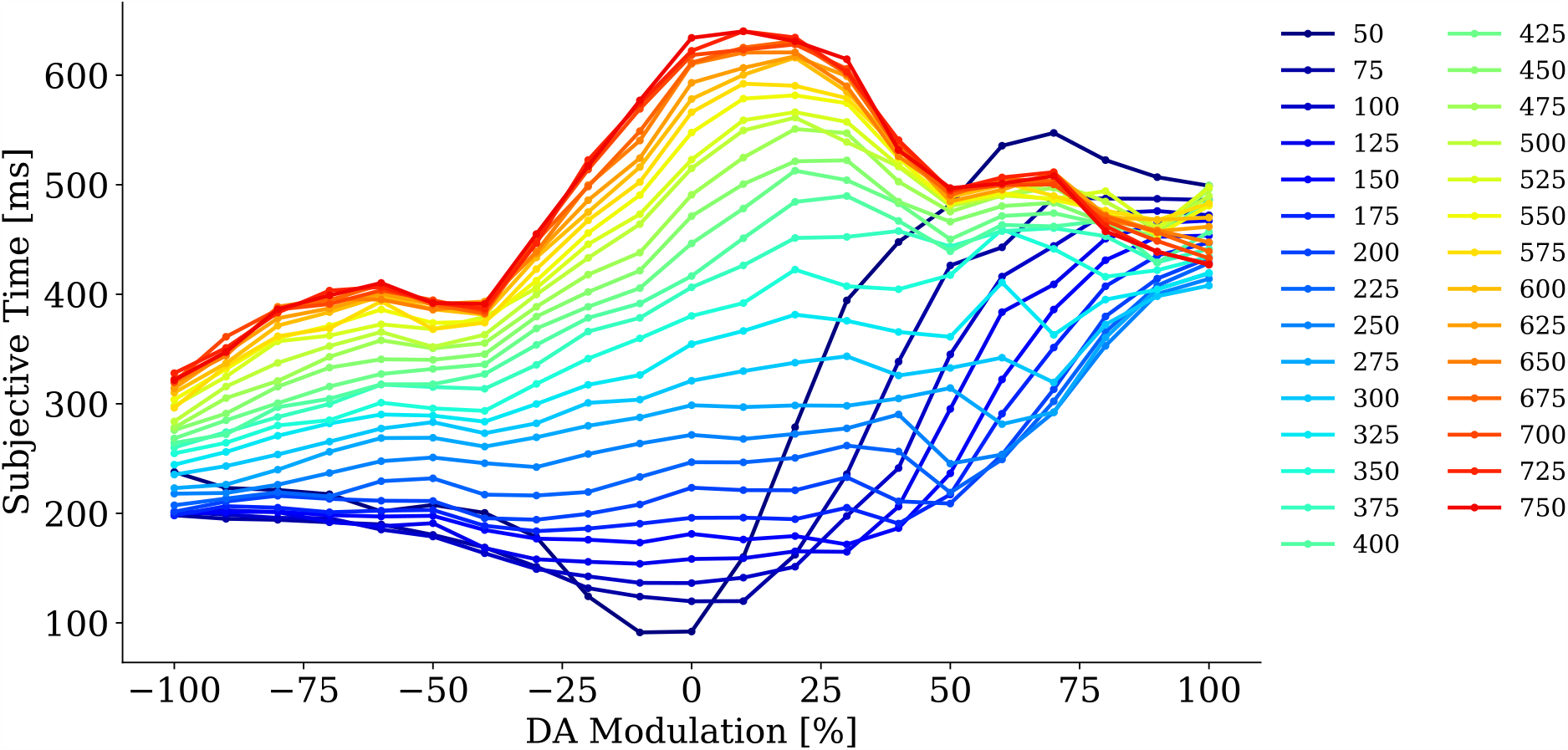
Dopamine modulation and the change of the subjective time within the full range. The estimated time for antagonistic (− 100 % - − 10 %) and agonistic (10 % - 100 %) dopaminergic modulation is presented here. Each color represents the tested interval, such that the evolution of the estimate of one interval can be considered over the modulation. An overestimation and an underestimation of time can only be found within the range of 200 - 600 ms and for percentages below ±50 %.

**SI Fig A.5:**
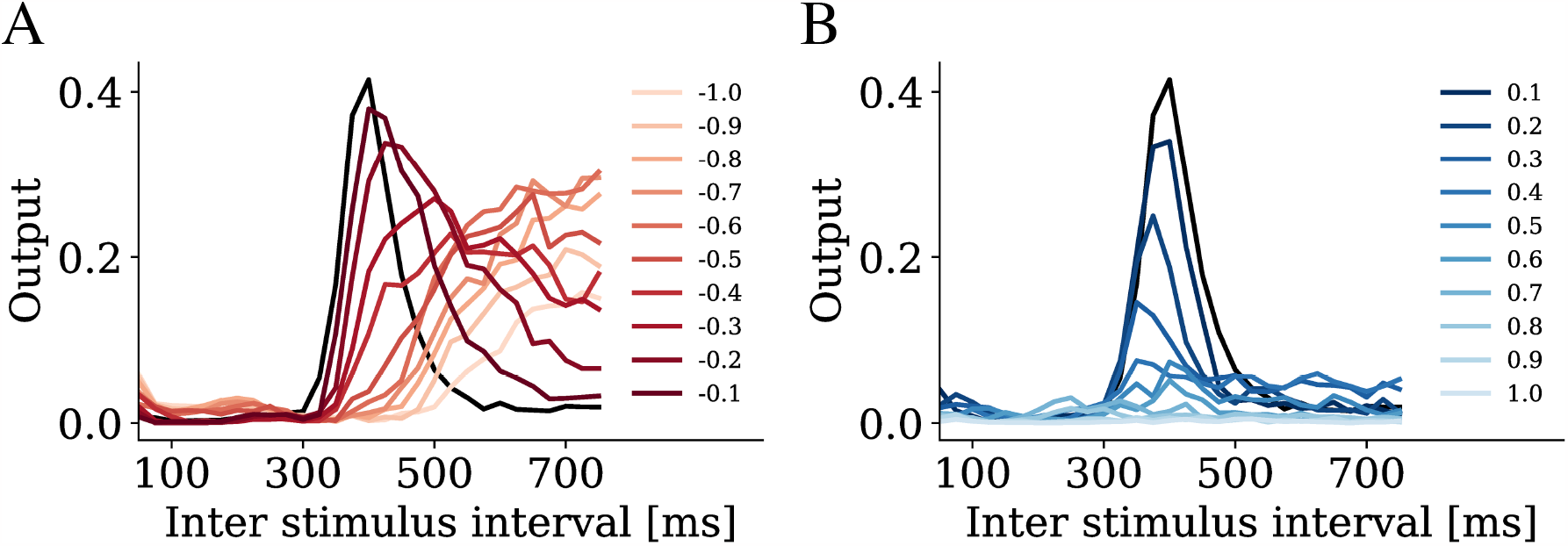
Outputs of the 400 ms readout neuron with altered DA modulation. **A** With increasing antagonistic dopaminergic modulation (from darker to lighter colors), the activation of the 400 ms interval-encoding unit is activated at later time points. **B** With increasing agonistic modulation the same unit is activated at earlier intervals. The output values decay with increasing modulation.

**SI Fig A.6:**
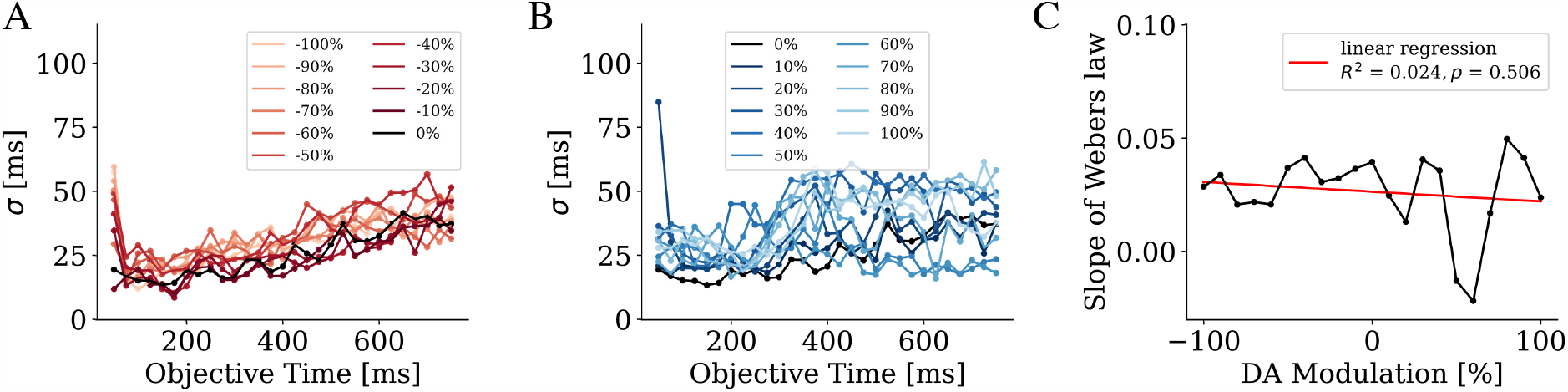
Standard deviations of the dopaminergic modulations. The standard deviations with respect to unmodulated Webers law (black) are shown for antagonistic **A** and agonistic **B** dopamine modulations of ±10 to ±100 % within the full range and **C** the slopes of the estimated times for each modulation fitted with a linear regression (red).

**SI Fig A.7:**
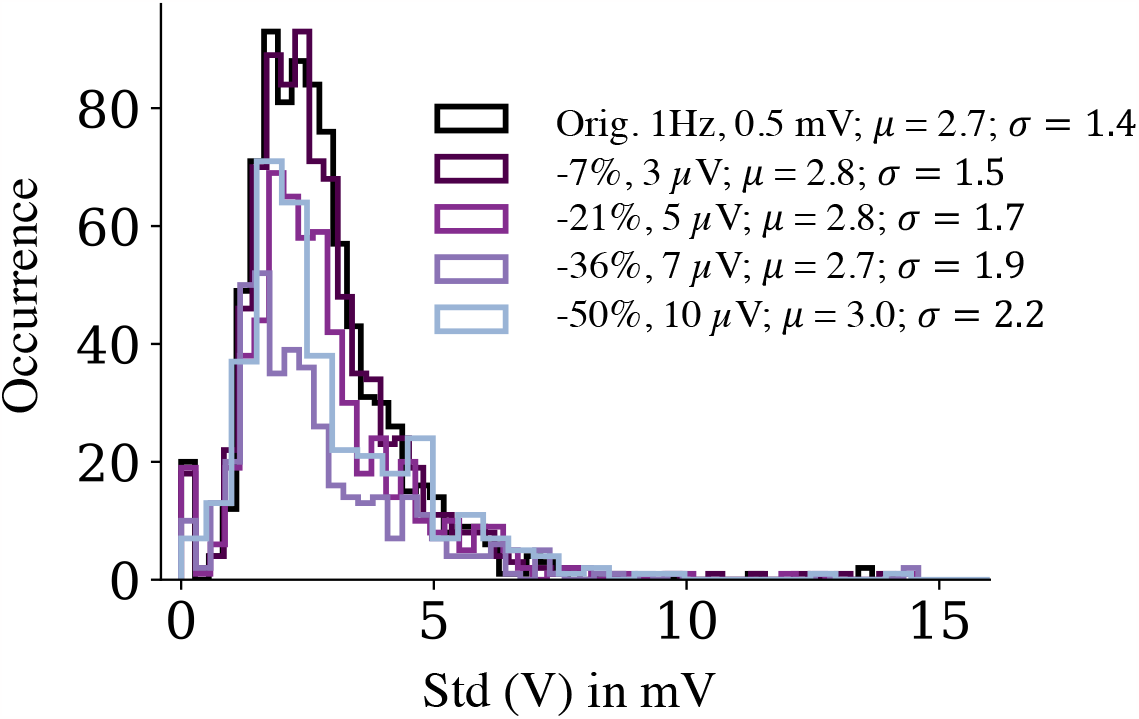
Subthreshold membrane potential fluctuations for different cases of Poisson noise. The standard deviation of the sub-threshold membrane potentials over time for non-zero firing neurons for the original simulation with 1 Hz in black and for different levels of Poisson noise with 1000 Hz. The mean *μ* and variance *σ* for the distribution of the histogram can be extracted from the legend.

**SI Fig A.8:**
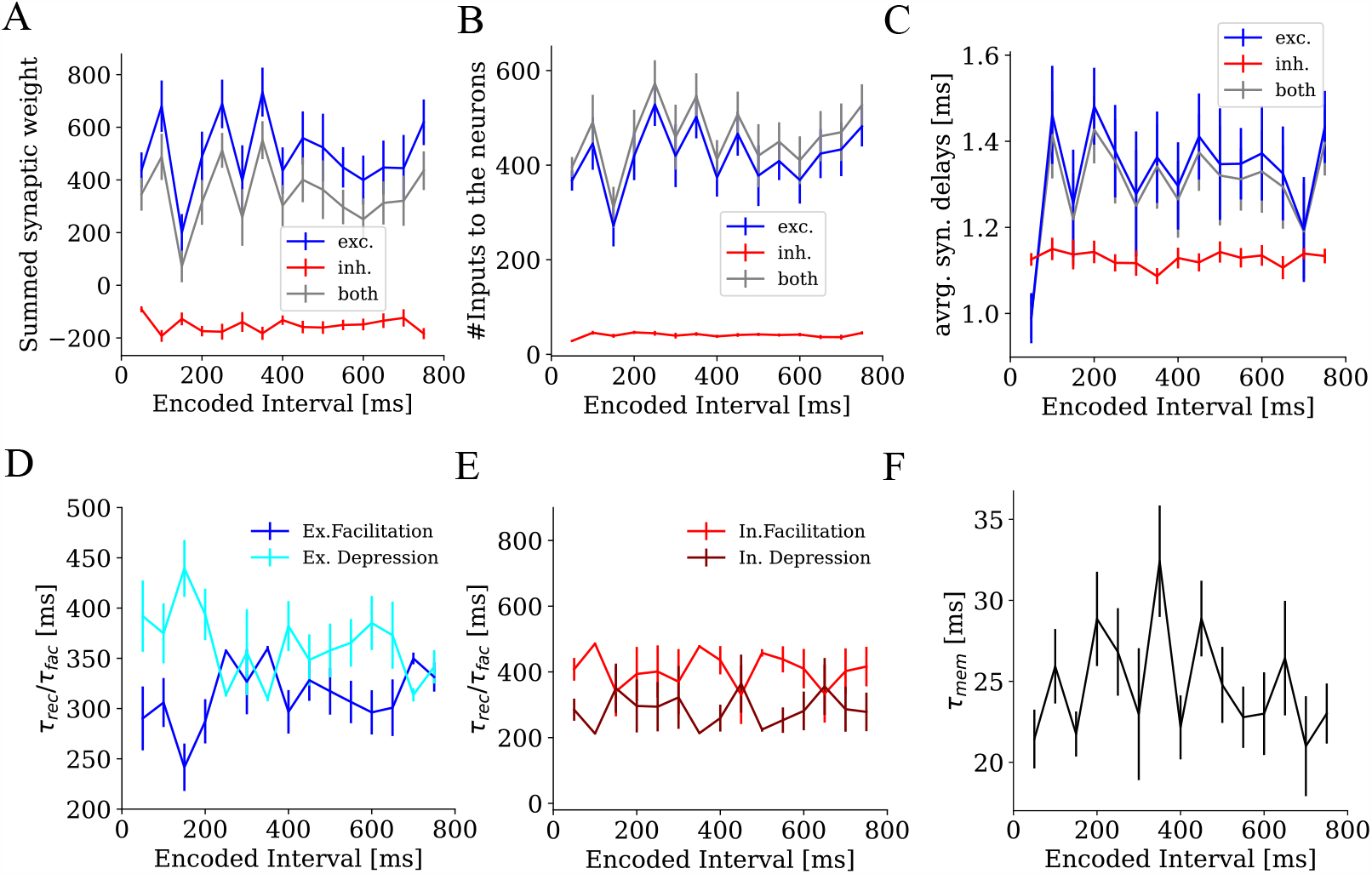
Neuronal and synaptic properties within interval-encoding pools below the readout threshold of 0.01. **A** The averaged summed synaptic weight **B**, the connectivities **C** and the averaged synaptic delays onto an interval-encoding pool for excitatory (blue), for inhibitory (red) and for all weights (black). For the synaptic delays, only inhibitory synaptic delays are significant (slope = −0.0002, *R*^2^ = 0.6, *p* = 0.02). The time constants of STP, *τ*_*fac*_ and *τ*_*rec*_, for excitatory **D** and inhibitory **E** neurons within a pool. **F** The averaged membrane time constants *τ*_mem_ for neurons within IEPs below the threshold for both neuron types.

**SI Table A.1:** Parameter adjustments for ablation analysis. The changes were applied in relation to the PFC model from [27].

| Ablation |  | Parameter changes<br>[%] |
| --- | --- | --- |
| **State-dependent PFC model** | $I_{\text{back}}$ | 1.40 |
| | $g_{\text{max,GABAB}}$ | 0.30 |
| | $g_{\text{max,GABAA}}$ | 0.30 |
| **Synaptic mechanisms** |  |  |
| no NMDA | $I_{\text{back}}$ | 140 |
| | $g_{\text{max,GABAB}}$ | 30 |
| | $g_{\text{max,GABAA}}$ | 30 |
| | $g_{\text{max,NMDA}}$ | 0 |
| no GABA<sub>B</sub> | - |  |
| no STP | $I_{\text{back}}$ | 430 |
| | $I_{\text{back,exc,layer 23}}$ | 130 |
| | $w_{\text{syn, exc}}$ | 80 |
| | $w_{\text{syn, inh}}$ | 40 |
| | $I_{\text{back,exc,layer 5}}$ | 210 |
|  | Synapse type | *staticsynapse (NEST)* |
| no NMDA, no GABA<sub>B</sub> & no STP | $I_{\text{back}}$ | 210 |
| | $I_{\text{back,exc,layer 23}}$ | 130 |
| | $w_{\text{syn, ex}}$ | 80 |
| | $w_{\text{syn, inh}}$ | 40 |
| | $I_{\text{back,exc,layer 5}}$ | 210 |
|  | Synapse type | *staticsynapse (NEST)* |
| | $g_{\text{max,GABAB}}$ | 0 |
| | $g_{\text{max,GABAA}}$ | 100 |
| | $g_{\text{max,NMDA}}$ | 0 |
| **Heterogeneity** |  |  |
| homogen. neurons | $I_{\text{back}}$ | 140 |
| | $I_{\text{back,exc}}$ | 130 |
| homogen. synapses | $I_{\text{back}}$ | 140 |
| | $I_{\text{back,exc}}$ | 110 |
| homogen. neurons, synapses | $I_{\text{back}}$ | 140 |
| | $I_{\text{back,exc}}$ | 120 |
| homogen. neurons, syn., 1 type of STP | $I_{\text{back}}$ | 140 |
| | $I_{\text{back,exc,layer 5}}$ | 220 |
| | $I_{\text{back,exc,layer 23}}$ | 180 |
| | $w_{\text{syn, inh}}$ | 280 |
| | $w_{\text{syn}}$ | 180 |
| homog. syn. delays | $\tau_D$ | 0.1 ms |
| **Irregular background activity** |  |  |

**SI Table A.2:** Linear regression was calculated for parameters (see Fig. 11) within the interval selective pools over the full range (50 - 750 ms).

| Type | linear regression |  |  |
| --- | --- | --- | --- |
| | slope | $p$ | $R^2$ |
| Exc. syn. weights $w$ | 0.75 | 0.004 | 0.47 |
| Inh. syn. weights $w$ | -0.18 | < 0.001 | 0.65 |
| Exc. inputs to the neurons $p_{\text{conn}}$ | 0.22 | 0.04 | 0.30 |
| Inh. inputs to the neurons $p_{\text{conn}}$ | 0.03 | < 0.001 | 0.65 |
| Exc. syn. delays $\tau_D$ | 0.0006 | < 0.001 | 0.65 |
| Inh. syn. delays $\tau_D$ | 0.00003 | 0.40 | 0.05 |
| Exc. facilitating time constants $\tau_{\text{fac,ex}}$ | 0.09 | 0.01 | 0.40 |
| Exc. depressing time constants $\tau_{\text{rec,ex}}$ | -0.11 | 0.01 | 0.42 |
| Inh. facilitating time constants $\tau_{\text{fac,in}}$ | 0.10 | 0.68 | 0.01 |
| Inh. depressing time constants $\tau_{\text{rec,in}}$ | -0.39 | 0.12 | 0.17 |
| $\tau_{\text{mem}} = C_m/g_L$ | 0.008 | 0.07 | 0.23 |

**SI Table A.3:** Ablation studies to identify the components of linear timing.

| Ablation | Slopes | Two sample<br>t-test of slopes<br>to original | IP<br>[ms] |
| --- | --- | --- | --- |
| **original** | 0.84 ±<br>0.01 | | 538.9 ±<br>17.1 |
| **Synaptic mechanisms** |  |  |  |
| no NMDA | 0.85 ±<br>0.01 | $t(497)=2.32$,<br>$p < .02$ | 532.1 ±<br>14.0 |
| no GABA<sub>B</sub> | 0.48 ±<br>0.03 | $t(396)=-56.89$,<br>$p < .001$ | 445.7 ±<br>14.0 |
| no STP | 0.61 ±<br>0.06 | $t(312)=-25.0$,<br>$p < .001$ | 380.5 ±<br>41.2 |
| no NMDA &<br>no GABA<sub>B</sub> & no STP | 0.35 ±<br>0.04 | $t(355)=-66.4$,<br>$p < .001$ | 424.1 ±<br>17.6 |
| **Heterogeneity** |  |  |  |
| homogen. neurons | 0.81 ±<br>0.03 | $t(491)=-5.92$,<br>$p < .001$ | 538.6 ±<br>7.8 |
| homogen. synapses | 0.86 ±<br>0.01 | $t(484)=-4.49$,<br>$p < .001$ | 548.2 ±<br>25.1 |
| homogen. neurons &<br>synapses | 0.84 ±<br>0.03 | $t(492)=-1.6$,<br>$p = 0.1$ | 406.8 ±<br>171.5 |
| homogen. neurons, syn.,<br>1 type of STP | 0.53 ±<br>0.08 | $t(304)=-31.34$,<br>$p < .001$ | 427.9<br>17.2 |
| homog. syn. delays | 0.86 ±<br>0.01 | $t(494)=4.03$,<br>$p < .001$ | 559.2 ±<br>19.0 |
| **Irregular background**<br>**activity** |  |  |  |
| no background<br>current | 0.71 ±<br>0.01 | $t(413)=-21.67$,<br>$p < 0.001$ | 390.5 ±<br>53.0 |

**SI Table A.4:** Ablation studies to identify the components of Weber’s law.

| Ablation | piecewise linear fit |  |  |
| --- | --- | --- | --- |
| | $a$ | $b$ | $c$ |
| **original** | 0.05 ±<br>0.01 | 9.4±<br>1.4 | 532.4 ±<br>75.1 |
| **Synaptic mechanisms** |  |  |  |
| no NMDA | 0.05 ±<br>0.004 | 9.5±<br>1.5 | 550.6±<br>35.8 |
| no GABA<sub>B</sub> | 0.05±<br>0.04 | 47.3±<br>9.0 | 663.7 ±<br>344.5 |
| no STP | 0.09±<br>0.03 | 29.4 ±<br>2.3 | 439.9±<br>138.1 |
| no NMDA, no GABA<sub>B</sub> & no STP | 0.03±<br>0.04 | 70.6±<br>11.0 | 429.0±<br>335.5 |
| **Heterogeneity** |  |  |  |
| homogen. neurons | 0.07 ±<br>0.04 | 14.5 ±<br>5.4 | 490.4±<br>182.2 |
| homogen. synapses | 0.04 ±<br>0.01 | 13.4 ±<br>2.9 | 645.1±<br>69.4 |
| homogen. neurons, synapses | 0.04 ±<br>0.01 | 15.2±<br>6.5 | 698.0 ±<br>239.9 |
| homogen. neurons, syn., 1 type of STP | 0.15±<br>0.17 | 48.6±<br>33.8 | 318.7±<br>376.1 |
| homog. syn. delays | 0.05 ±<br>0.001 | 10.8 ±<br>2.7 | 540.6 ±<br>82.0 |
| **Irregular background activity** |  |  |  |
| no background current | -0.07±<br>0.1 | -8.8 ±<br>112.9 | 658.9±<br>900.9 |

## Appendix B. Supplementary materials S2

### Origin of the scalar property for more than one pool and output neuron

In the main text, we have shown how the scalar property arises from the stereotypical firing rate profiles across interval-encoding pools IEPs and the fixed relation between the mean and the standard deviation of the firing rates according to the binomial distribution. We only considered the special case that a) the estimated interval is one of the intervals that are encoded in one of the pools (*t* = *T*_*i*_), and b) there is only a single pool and output neuron, both of which encode this interval *T*_*i*_. While this case is instructive to understand the mechanism of the scalar property, it is necessary to generalize the considerations of the main text to arbitrary interval durations and multiple pools and output neurons (although usually, a few pools and output neurons are sufficient due to the narrow tuning curves). A full mathematical derivation of this case is beyond the scope of the current paper. Instead, we outline the general rationale here and aim to verify each of the corresponding assumptions with data from the network model.

SI Fig. B.9A shows the relation between the mean and the standard deviation of the activity of each of the output neurons *O*_*j*_ (cf. lines and shaded areas in Fig. 2A). This relation can be well-fitted to the function 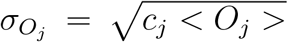 with a relatively constant *c*_*j*_ across output neurons *j* (0.26 ± 0.04). As the duration estimate *T*_est_(*t*) is formed by summing over the activities *O*_*j*_ of each output neuron multiplied by the duration *T*_*j*_ that neuron represents, mean and standard deviation of the estimate are given by 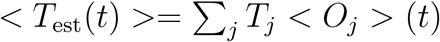 and 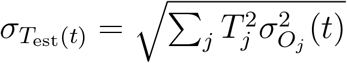, respectively.

Assume that we want to compute both quantities at a given time *t*^∗^. Under the assumption that all tuning curves *< O*_*j*_ *>* have the same shape, namely mean-shifted Gaussians with constant standard deviations (SI Fig. B.10A), we can interpret the values *< O*_*j*_(*t*^∗^) *>* of the different output functions *j* at this time *t*^∗^ as the values of a single function *< O*(*t*) *>* at different times *t* that reflect the shift of the mean of each of the different Gaussians (SI Fig. B.10B). Given the fact that the output activities are normalized such that Σ_*j*_ *< O*_*j*_(*t*) *>*= 1 for all times *t*, we can interpret *< O*(*t*) *>* as the probability distribution of the time *t* elapsed relative to *t*^∗^. Thus, the average of elapsed time with respect to *< O*(*t*) *>* (or *< O*_*j*_(*t*^∗^) *>*, equivalently) is *t*^∗^ and we can write

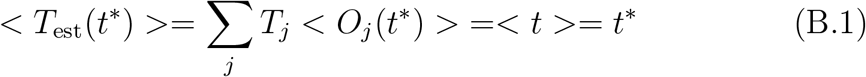

**SI Fig A.9:**
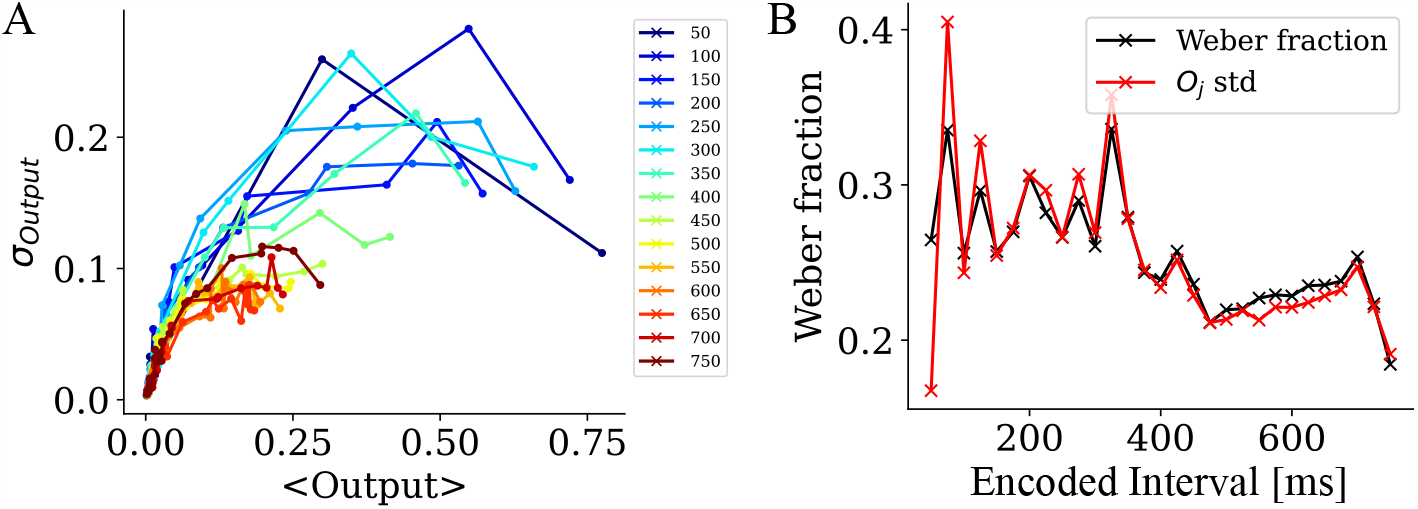
Assessing the origin of the scalar property. A.Standard deviations of the output neuron activity as a function of their mean activity. B.Weber fractions (black curve, standard deviation of the duration estimated divided by its mean) and square root of the sum of the variances of all output neurons (red curve) for each of the encoded interval durations.

and, using 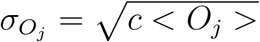 (see above) and Var(*t*) =*< t*^*2*^ *>* − *< t >*^*2*^,

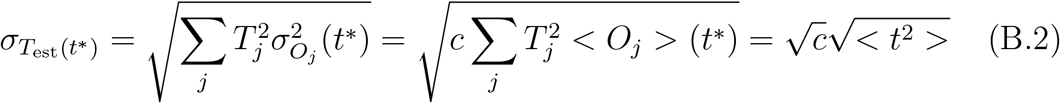

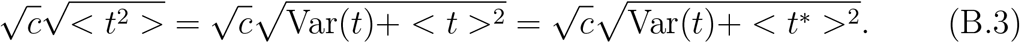

Var(*t*) is a constant that reflects the width of the tuning curves (about 25 ms for the almost constant curves representing interval durations up to 300 ms) while 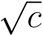is the Weber fraction.

Together, eqn. B.1, B.2 and B.3 yield the generalized Weber law [53]

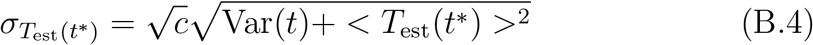

In the generalized Weber law, a constant error dominates the standard deviations of the timing estimates from short durations, resulting in a decreasing Weber fraction, until the linear part dominates for longer intervals and the Weber fraction becomes constant. Indeed, the state-dependent PFC model shows this often-observed behavior of the Weber fraction (Fig. 2D).

We test the accuracy of the above calculations by computing the Weber fraction 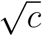 from the data in two ways: First, as the ratio between the standard deviations 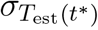 and the means *< T*_est_(*t*^∗^) *>*, which yields 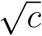 for long enough intervals (see above). And second, as the square root of the sum of the variances in all output ne urons *O*_*j*_. This second relation, 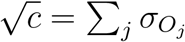 follows from the normalization Σ_*j*_ *< O*_*j*_(*t*) *>*= 1 in combination with 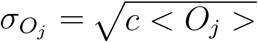 . Indeed, the results of the two calculations are in good agreement with each other (SI Fig. B.9B), except for the expected deviations for short intervals.

**SI Fig A.10:**
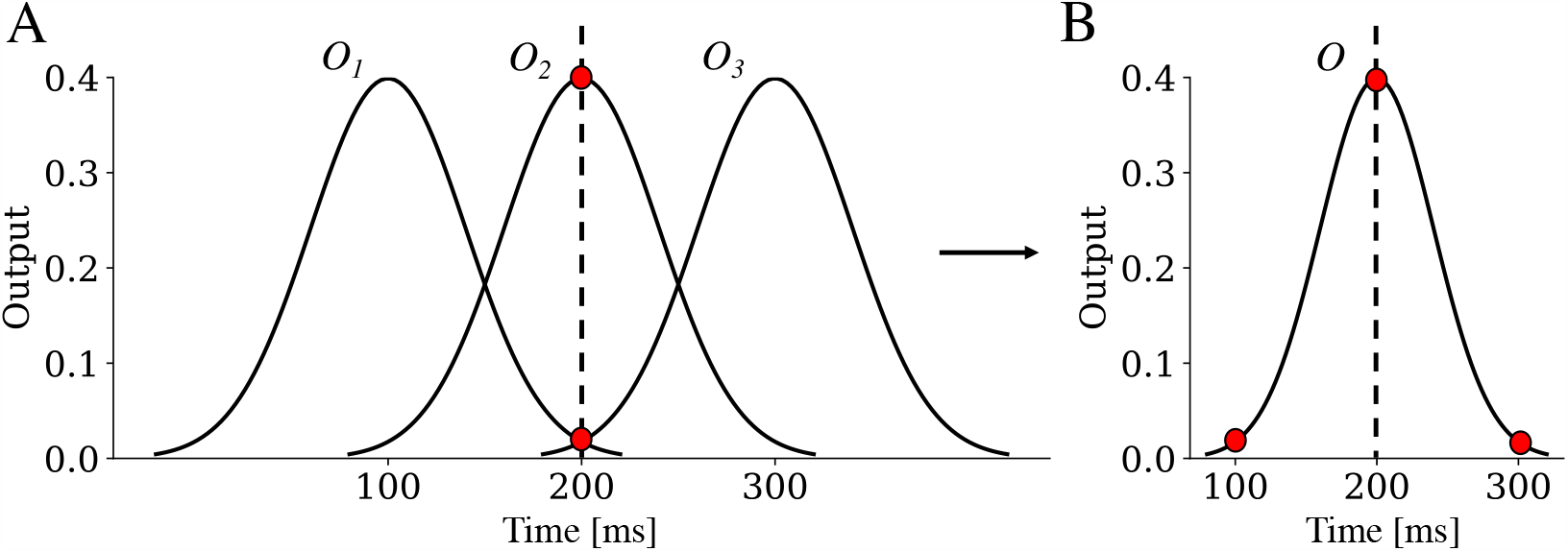
Interpreting the tuning curves of the output neurons as a probability density function. **A** Illustration of multiple tuning curves (mean-shifted Gaussians with constant standard deviations) evaluated at a given time *t*∗ . **B** The values of the different tuning curves *O*_*j*_ can be interpreted as the values of a single Gaussian *O* (centered at *t*∗) at different times that correspond to the shifted means. Note that the area under this Gaussian is one, so *O*(*t*) can be interpreted as a probability distribution function for the time *t* elapsed relative to *t*∗.

